# Differential Expression of TKS4 Isoforms and Their Role in Cellular Processes in Breast Cancer

**DOI:** 10.64898/2026.07.22.740038

**Authors:** Serhii Kropyvko, Nazar Shevchuk, Olga Gubar, Kyrylo Lavrynenko, Yaroslav Nemesh, Denys Kozakov, Valentyn Polishchuk, Valentyna Kryklyva, Liubov Syvak, Nataliia Verovkina, Tetyana Gryaznova

**Affiliations:** Department of Functional Genomics, Institute of Molecular Biology and Genetics NAS of Ukraine, Kyiv 03143, Ukraine; State Institution National Research Center for Radiation Medicine of National Academy of Medical Sciences of Ukraine, Kyiv, Ukraine

**Keywords:** TKS4 isoforms, gene expression, MCF-7 cells, breast cancer, epithelial-mesenchymal transition, invasion, migration, proliferation

## Abstract

The scaffold protein TKS4 plays a role in the development of several cancers. Alternative splicing of the *TKS4* gene generates two isoforms, TKS4L and TKS4b; however, their distinct expression patterns and functional roles have not yet been characterized. We have shown that TKS4 isoforms were differentially expressed across human cell lines and breast cancer (BC) tumor samples. Both *TKS4L* and *TKS4L*/*TKS4b* mRNA ratios were significantly altered in tumors compared with adjacent tissues. We identified six novel binding SH3-domain-containing partners for TKS4L, none of which interact with TKS4b, suggesting their functional differences. Tyrosine phosphorylation of both isoforms was induced by Src(Y527F) kinase overexpression, enabling binding to the SH2 domains of signaling proteins. Interestingly, TKS4b significantly accumulated in the nucleus, while TKS4L was primarily present in the cytosol in MCF-7 cells. TKS4b overexpression enhanced MCF-7 cell migration. Both TKS4 isoforms exhibit oncogenic properties by promoting epithelial-mesenchymal transition in BC cells, highlighting their potential as targets for therapeutic intervention.

## INTRODUCTION

The scaffold protein TKS4 (tyrosine kinase substrate with four SRC homology 3 (SH3) domains), encoded by the *SH3PXD2B* gene, belongs to the family of tyrosine kinase substrates that also includes the structurally and functionally related protein TKS5. TKS4 comprises an N-terminal PX domain that interacts with membrane phosphatidylinositol phosphates (PtdInsPs), four SH3 domains, and multiple proline-rich motifs that mediate binding to partner proteins (Figure 1).

**Figure 1.**
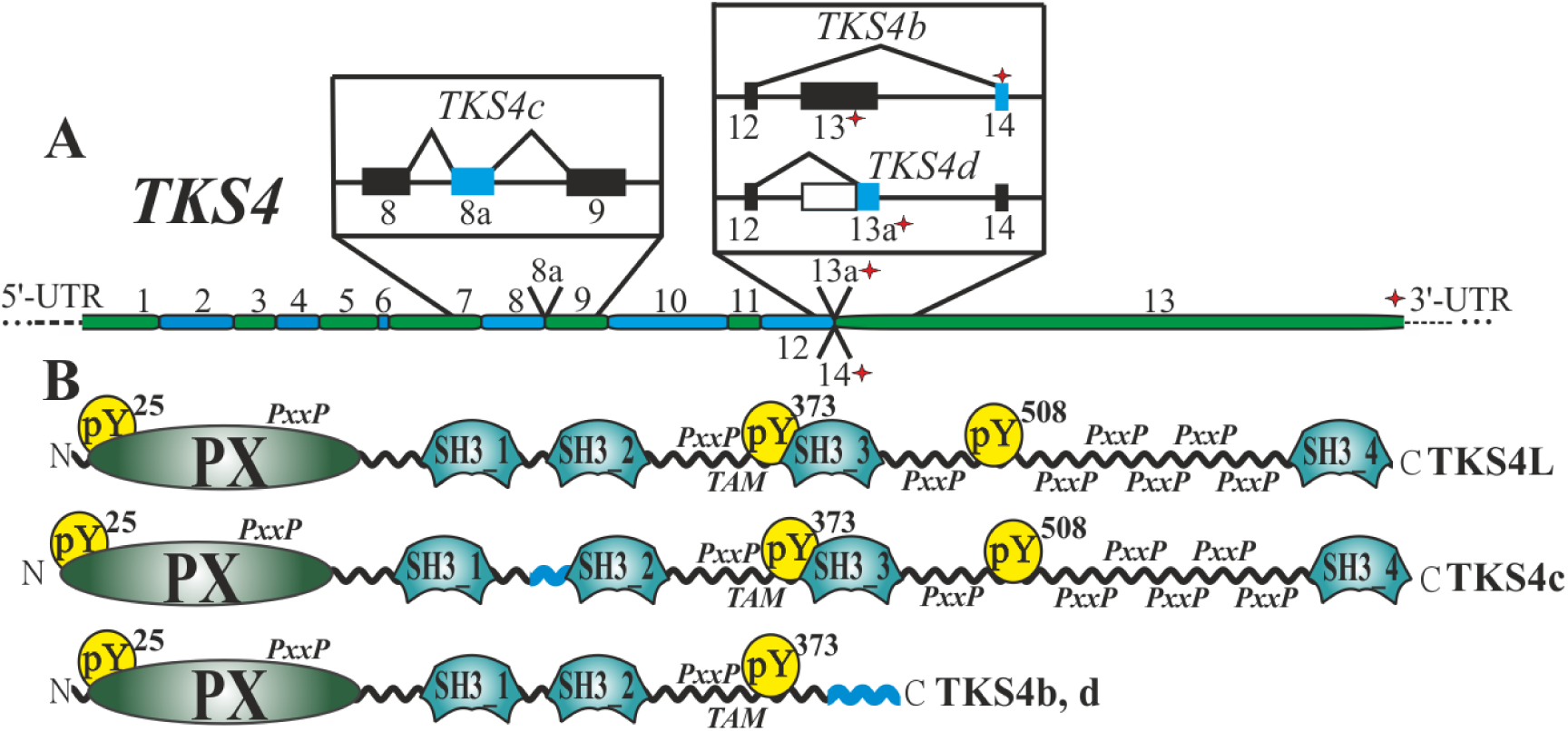
**TKS4 isoforms**. **(A)** Schematic representation of the *TKS4* gene mRNA and alternative splicing events leading to the formation of the *TKS4b*, *TKS4c* and *TKS4d* transcripts. **(B)** Domain structure of the TKS4L, TKS4b, TKS4c and TKS4d protein isoforms. pY^n^ – tyrosine phosphorylation sites; PxxP – proline-rich motifs; TAM – tandem SH3-associated motif responsible for the closed autoinhibitory conformation of TKS4; the additional sequence at the C-terminus of the TKS4b/TKS4d and coding 8a exon of the TKS4c isoforms are marked in blue.

TKS4 is regulated by SRC kinase–dependent phosphorylation of tyrosine residues Y25, Y373, and Y508, a modification that is critical for its role in intracellular signaling and subcellular localization.^1^ As a scaffold protein, TKS4 facilitates the assembly and coordination of signaling complexes by simultaneously engaging multiple interacting partners.^1,2^

Functionally, TKS4 is implicated in cell motility and invasion, the regulation of actin polymerization, and the formation of membrane protrusions, including podosomes, membrane ruffles, and related structures.^3^ It is a key regulator of invadopodia maturation, actin-rich membrane structures that support malignant cell migration and metastatic dissemination.^4^ TKS4 also contributes to the secretion of matrix metalloproteinases,^1,5^ adipogenesis and osteogenesis,^6,7^ angiogenesis,^8^ epithelial–mesenchymal transition (EMT),^9,10^ and Nox1-dependent reactive oxygen species production.^11^ Moreover, TKS4 is secreted in exosomes in an MT1-MMP–dependent manner, thereby transferring invasive properties to MDA-MB-231 cells.^5^ Consistent with these functions, TKS4 is widely regarded as a marker of invadopodia^4^ and of adipose stem and progenitor cells.^7^

Dysregulation of *TKS4* is associated with several pathological conditions, including Frank– Ter Haar syndrome^12^ and steatotic liver disease.^7^ Altered *TKS4* expression has been reported in lung cancer,^9^ melanoma,^13^ hepatocellular carcinoma,^14^ oral squamous cell carcinoma,^15^ and colorectal cancer.^16^ However, these studies did not address potential differences in the expression of distinct *TKS4* isoforms generated by alternative mRNA splicing.

To date, functional investigations have generally not distinguished between TKS4 isoforms, with the majority of existing data attributed to the protein without regard to its spliced variants. In contrast, GenBank annotations identify two transcripts generated by alternative splicing of the terminal exon of *SH3PXD2B*: the canonical *TKS4L* and an additional isoform, *TKS4b*. Previous studies have examined the *SH3PXD2B* gene function at the locus level without distinguishing isoform-specific contributions, including expression analyses using TCGA and GENT2 by Tilajka et al.,^16^ knockout approaches by László et al.,^9^ CRISPR/Cas9-mediated knockout by Szeder et al.,^10^ and qPCR-based expression analyses by Kui et al.^14^ and Hubiernatorova et al.^17^ Wenzel et al.^5^ employed siRNAs specifically targeting *TKS4L*, whereas Iizuka et al.^13^ and Mehes et al.^8^ did not specify the nucleic acid sequences used, leaving the isoform specificity of their analyses unresolved.

Furthermore, many antibody-based studies have historically focused on TKS4L, as signals obtained with antibodies targeting N-terminal regions shared among multiple variants were often attributed exclusively to TKS4L, without considering potential contributions from other isoforms (Table S1). In the present study, we use both pan-TKS4 antibodies recognizing epitopes common to all variants and antibodies directed against isoform-specific sequences.

In the present study, we characterize the expression patterns of *TKS4* isoforms across a panel of cell lines and human breast cancer (BC) specimens, and further investigate their functional roles in the MCF-7 cell line. BC is a highly heterogeneous malignancy, representing the most frequently diagnosed cancer in women and a major cause of cancer-related mortality globally. According to the International Agency for Research on Cancer (IARC), in 2022 BC ranked second in overall incidence (11.5%) and first among women (23.8%), with approximately 2.2 million new cases and a mortality rate of 6.8%, corresponding to more than 600,000 deaths.^18^ Despite substantial therapeutic progress, a considerable proportion of patients develop relapse or metastatic disease due to intrinsic or acquired therapy resistance, underscoring the need for additional strategies to improve long-term outcomes.^19^ Previous studies in BC models have demonstrated the critical role of TKS4 in the formation of various types of invadopodia and, consequently, in tumor metastasis.^20,21^ However, the contribution of distinct TKS4 isoforms to other processes relevant to BC progression, including EMT, cell proliferation etc., has not been investigated. TKS4 has also been shown to regulate signaling downstream of tyrosine kinase receptors such as EGFR, which are frequently dysregulated in BC. Investigating the specific functions of TKS4 isoforms may therefore provide deeper insight into the molecular mechanisms of BC development and metastasis. A comprehensive analysis of the TKS4 interactome and isoform-resolved expression profiles in normal and malignant breast tissues is expected to provide deeper insight into the contribution of scaffold protein isoforms to tumor initiation and progression and to clarify their utility as prognostic biomarkers in cancer.

## RESULTS

### Alternative splicing of *TKS4* mRNA

The human *TKS4* gene (*SH3PXD2B*; also referred to as *FTHS*, *HOFI*, *TSK4*, *FAD49*, and *KIAA1295*) is located on chromosome 5. According to GenBank annotations, this gene gives rise to three transcripts: the canonical long isoform of approximately ∼101 kDa (apparent molecular mass of ∼120 kDa by SDS-PAGE) (*TKS4*, *TKS4L*, *TKS4a*; NM_001017995.3) and alternative isoforms, *TKS4b* (NM_001308175.1), generated by exclusion of the 13th exon and inclusion of the 14th exon in the mature mRNA (Figure 1) and *TKS4c* (NM_001445599.1) generated by inclusion of the 8a exon in the mature mRNA. The TKS4b isoform has a predicted molecular mass of ∼49 kDa and comprises the N-terminal PX domain, the first two SH3 domains (SH3_1 and SH3_2), and a unique C-terminal extension of 34 amino acid residues (DHKNVHLESPVEVPLRRIKMRTALGKHVAASMEM). The TKS4c isoform has a predicted molecular mass of ∼105 kDa, comprises all domains of the long isoform and extension of 28 amino acid residues (VSWRYWSLPRPVGRRRTLGDLYAISWRQ) between domains SH3_1 and SH3_2.

An additional alternatively spliced transcript is annotated in the Ensembl Genome Browser (ENST00000518522_5). This variant is produced via utilization of an internal splice acceptor site within exon 13, resulting in the removal of most of this exon as an intron (6147 nucleotides). In the present work, this transcript is designated as *TKS4d*. The *TKS4d* transcript is predicted to encode a protein isoform of similar molecular size and an identical domain architecture to TKS4b, with an N-terminal PX domain and two SH3 domains, but harboring a distinct C-terminal tail composed of 14 unique amino acids (EQWALPTTSCLLIG; Figure 1).

### Expression of *TKS4* transcripts in human cell lines of various origins

Expression of *TKS4* transcripts was quantified by qPCR in human cell lines using isoform-specific primers for *TKS4L*, *TKS4b*, and *TKS4d*, as well as primers targeting exons 1–3, common to all known *TKS4* variants. The data revealed cell type–dependent heterogeneity in both individual isoforms and total *TKS4* transcript pools. *TKS4L* mRNA was detected in all analyzed cell lines, with highest levels in U-251 MG (glioblastoma), DU145 (prostate carcinoma), and SH-SY5Y (neuroblastoma), and lowest levels in HeLa (cervical carcinoma) and A431 (epidermoid carcinoma). *TKS4b* transcripts were absent in HeLa and A431 cells and most abundant in DU145 cells. *TKS4d* mRNA was detected in all cell lines except HeLa, but generally at much lower levels than *TKS4L* and *TKS4b*, with the exception of DU145 and KRC/Y, where *TKS4d* was relatively enriched. Total *TKS4* transcript levels largely mirrored isoform-specific expression and were detectable in all cell lines (Figure 2A). The relative proportions of *TKS4L*, *TKS4b*, and *TKS4d* varied markedly between cell types (Figure 2B–D, Table S2). The altered ratio between *TKS4* transcripts may indicate the presence of a regulatory mechanism that maintains the balance between alternatively spliced variants. Such shifts in isoform ratios are often associated with functional specialization of transcript variants and may influence the composition of protein complexes involved in invadopodia formation or other processes.

**Figure 2.**
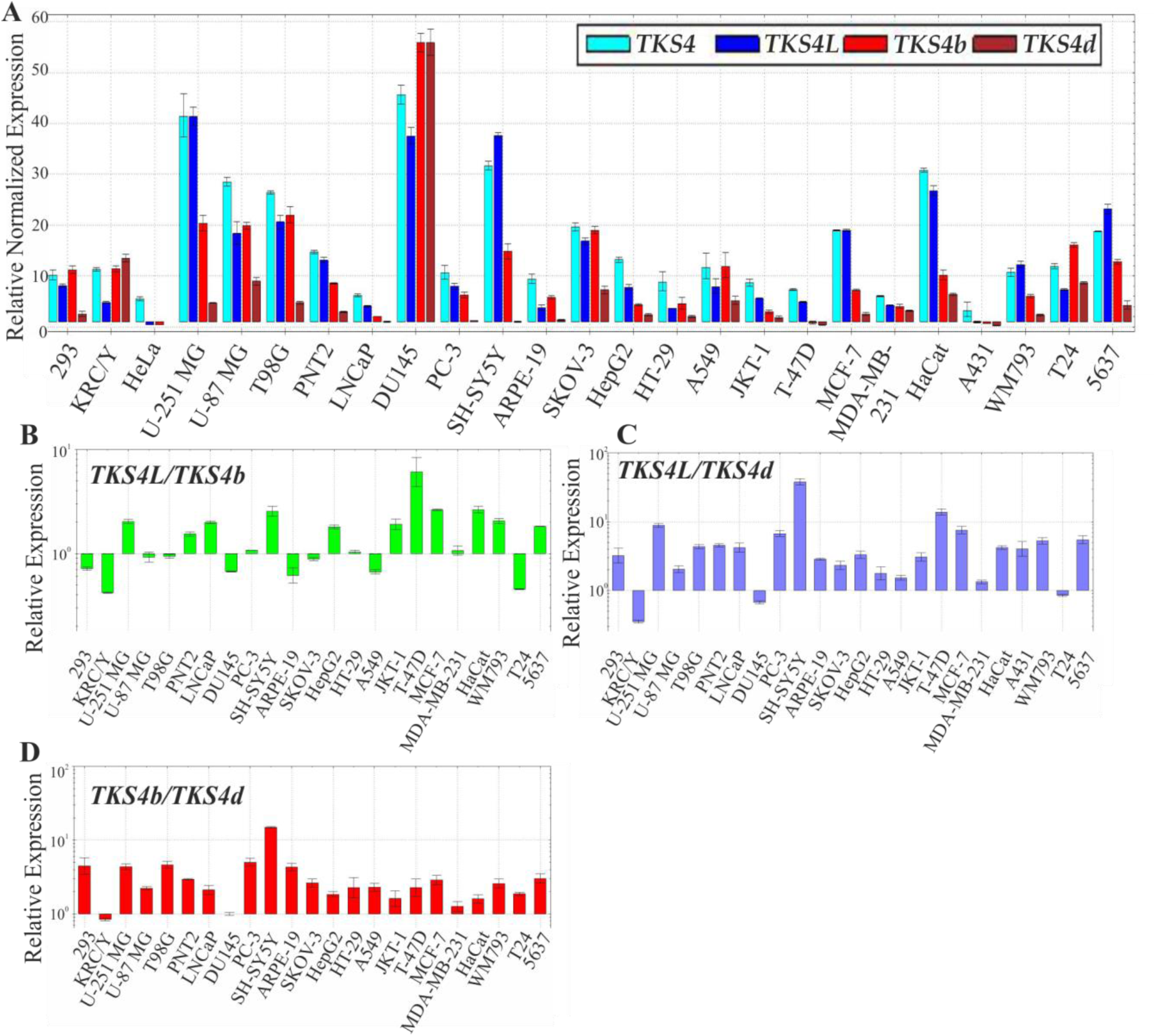
Expression of *TKS4* mRNA and protein isoforms in human cell lines of various origins. **(A)** Expression of *TKS4* total, *TKS4L*, *TKS4b*, and *TKS4d* transcripts; linear presentation of the data. Expression normalization to the *TBP* gene with standard error of the mean (lg). Ratio of *TKS4L*/*TKS4b* (**B**), *TKS4L*/*TKS4d* (**C**), and *TKS4b*/*TKS4d* (**D**) transcripts; Log10 presentation. Tukey’s HSD test was used.

### Expression profile of *TKS4* transcripts in breast tumors

The total *TKS4* mRNA pool has been proposed as a potential biomarker in several malignancies, including melanoma,^13^ lung cancer,^9^ hepatocellular carcinoma,^14^ and oral squamous cell carcinoma.^15^ We therefore examined whether *TKS4* transcript expression is altered in BC, one of the most prevalent and clinically aggressive tumor types.

Using the GENT2 database, which reports microarray-based mRNA expression, we compared *TKS4* expression between normal and tumor tissues across multiple cancer types. Depending on tumor origin, *TKS4* levels were higher, lower, or unchanged relative to normal tissue; BC belonged to the latter group, with no apparent difference in total *TKS4* transcript levels between tumors and controls (Figure 3A,S1; Table S3). A similar analysis using RNA-seq data from The Cancer Genome Atlas Breast Invasive Carcinoma (TCGA-BRCA) using the UCSC Xena browser cohort likewise revealed no significant differences in total *TKS4* expression between normal and tumor tissues or among BC subtypes (Figure 3B). Using the same analytical platform to evaluate the individual transcripts *TKS4L*, *TKS4b*, and *TKS4d*, no significant differences in expression were observed between normal breast tissues and breast tumors when the samples were analyzed without stratification by molecular subtype (Figure S2). In contrast, analysis of *TKS4L* expression in the TCGA-BRCA cohort using the SpliceSeq platform revealed lower expression in LumB HER2+ tumors than in LumA, Basal-like, and HER2-enriched subtypes, whereas, no significant difference was observed compared with Normal-like tissues (Figure 3C). Similar analyses could not be performed for *TKS4b* and *TKS4d* because these isoforms cannot be distinguished using the SpliceSeq platform.

**Figure 3.**
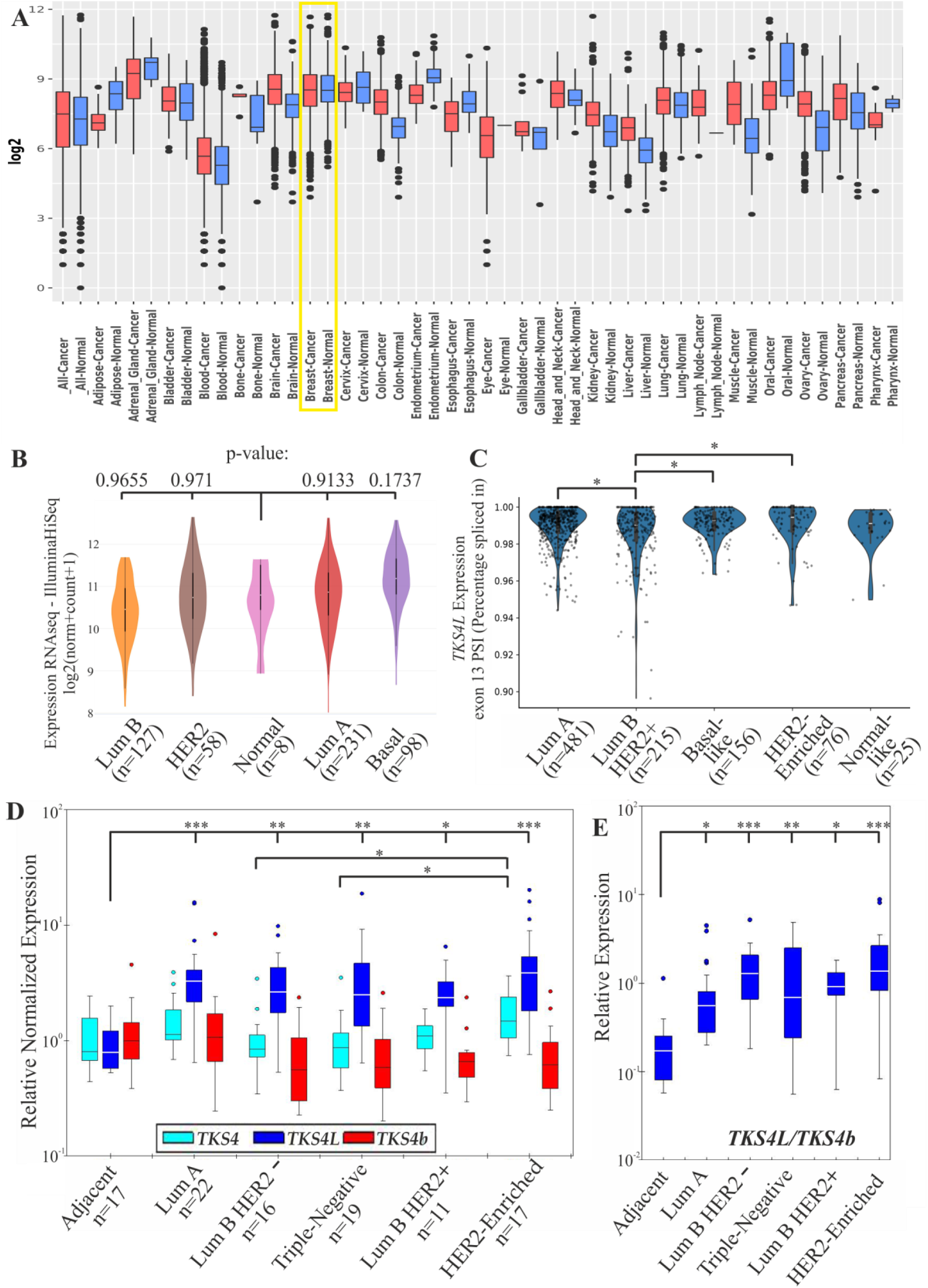
Expression of the *TKS4* gene in cancer. **(A)** *TKS4* expression in normal and tumor samples in various cancer types based on the GENT2 database. The BC (n=1246) is highlighted with a box (full version – Figure S1). **(B)** The TCGA database was analyzed via the UCSC Xena browser to compare *TKS4* expression in normal and tumor breast tissues; Data is presented as Log2 (norm+count+1)-transformed. **(C)** The TCGA database was analyzed via the SpliceSeq browser to compare *TKS4L* expression in normal and tumor breast tissues; post hoc Dunn test was used * – *p* < 0.05; data are presented as exon 13 Percentage spliced in (PSI). **(D)** Relative expression levels of the total pool of *TKS4* transcripts, *TKS4L*, and *TKS4b* in human BC samples from LumA (n=22), LumB HER2– (n=16), triple-negative (n=19), LumB HER2+ (n=11), and HER2-Enriched (n=17) tumor types compared with adjacent tissues (Adjacent, n=17), and (**E**) the ratio of *TKS4L*/*TKS4b* transcripts. Data is presented as Log10-transformed. The expression ΔΔCq was normalized to that of the *TBP* gene with the standard error of the mean (lg). One-way ANOVA * – *p* < 0.05, ** – *p* < 0.01, *** – *p* < 0.001.

To resolve isoform-specific patterns, we quantified *TKS4L*, *TKS4b*, and total *TKS4* mRNA by qPCR in RNA isolated from human BC specimens of different molecular subtypes and their matched adjacent tissues. Tumors were classified according to estrogen receptor (ER), progesterone receptor (PR), and HER2/neu status into luminal A (LumA), luminal B HER2-negative (LumB HER2–), triple-negative, luminal B HER2-positive (LumB HER2+), and HER2-enriched subtypes.^22^ *TKS4L* transcripts were significantly upregulated (2.6-4.1-fold) in all tumor types compared with adjacent tissues, with the highest levels observed in HER2-enriched tumors. In contrast, *TKS4b* transcript levels did not differ significantly between tumors and adjacent tissues, although a trend toward reduced expression in tumors was evident. Consistent with public datasets, the total *TKS4* mRNA pool did not differ between tumors and adjacent tissues; however, among BC subtypes, total *TKS4* expression was significantly lower in LumB HER2– and triple-negative tumors than in HER2-enriched tumors (Figure 3D; Table S4). Analysis of *TKS4L*/*TKS4b* ratios revealed marked isoform redistribution. In adjacent tissues, *TKS4b* predominated (ratio 0.21), and this pattern was preserved in LumA, triple-negative, and LumB HER2+ tumors (ratios 0.67, 0.89, and 0.78, respectively). By contrast, *TKS4L* was dominant in LumB HER2– (ratio 1.2) and HER2-enriched tumors (ratio 1.28). In all tumor subtypes, the *TKS4L*/*TKS4b* ratio differed significantly from that of adjacent tissue (Figure 3E; Table S5).

Collectively, these data indicate that only *TKS4L* mRNA is robustly upregulated in BC relative to adjacent tissue, and that isoform balance is shifted toward *TKS4L* in specific subtypes. Thus, *TKS4L* expression levels and/or the *TKS4L*/*TKS4b* ratio may serve as candidate markers of malignancy and molecular subtype in human breast cancer.

### Protein partner associations of TKS4L and TKS4b isoforms

A defining feature of the scaffold protein TKS4 is its multiple proline-rich motifs, which mediate interactions with SH3 domain–containing partners. A long linker between the third and fourth SH3 domains harbors eight proline-rich motifs, and an additional motif is located in the linker between the second and third SH3 domains. A further PxxP motif within the PX domain suggests that this domain may also participate in SH3-mediated interactions.^23^

To identify potential isoform-specific association partners of TKS4L and TKS4b, we performed GST pull-down assays using GST-fused SH3 domains from the following proteins: the endocytic factors AMPH1 and BIN1; the adaptor proteins NCK1, NCK2, CRK, and TKS5; phospholipase PLCγ1; the membrane-deforming proteins IRSp53 and IRTKS; and the kinase CSK. For in vitro binding assays, we used a stable T-REx-293 cell line overexpressing omni-tagged TKS4b, and detected endogenous TKS4L and omni-TKS4b with anti-SH3PXD2B and anti-omni antibodies, respectively. Endogenous TKS4L efficiently precipitated with AMPH1, BIN1, CRKI, PLCγ1 (Figure 4A), SH3(2) domains of NCK1 and NCK2 (Figure 4B), and the fifth SH3 domain of TKS5 (SH3(5); Figure 4C). In contrast, no interactions of omni-TKS4b with these SH3 domains were detected (Figure 4A–C).

**Figure 4.**
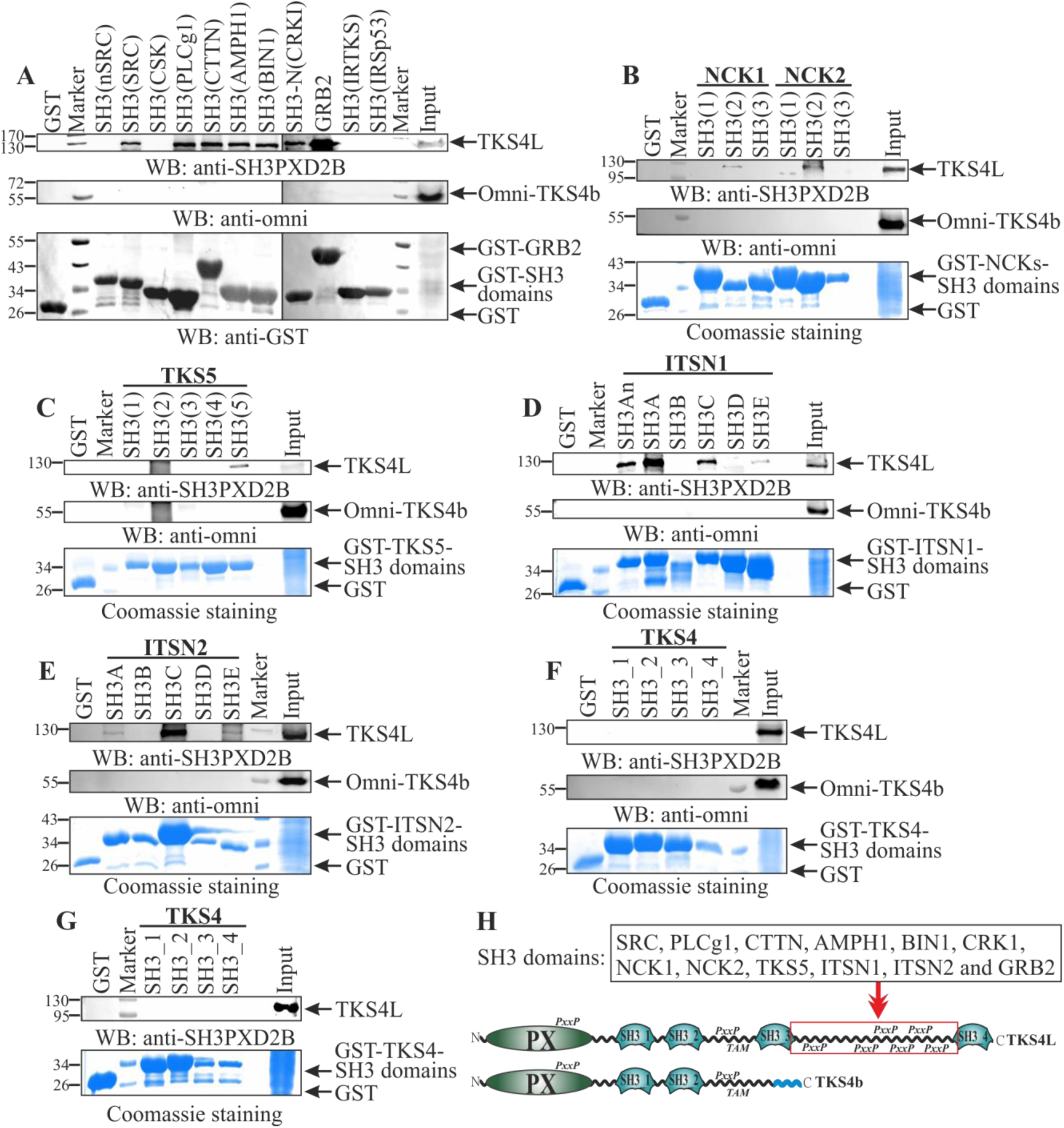
TKS4L and TKS4b isoforms associate with SH3-containing proteins. The GST-SH3 domains of SRC (neuronal and ubiquitously expressed), CSK, PLCγ1, CTTN, AMPH1, BIN1, CRK, GRB2, IRTKS, IRSp53 (**A**), NCK1, NCK2 (**B**), TKS5 (**C**), ITSN1 (**D**), ITSN2 (**E**), and TKS4 (**F-G**) were expressed in bacteria and affinity purified. The GST-SH3 domains and GST alone (control), which were immobilized on glutathione beads, were incubated with lysates from T-REx-293 cells with or without expression inducible omni-TKS4b. Bound proteins were separated via SDS-PAGE and detected by immunoblotting via antibodies against TKS4L (Anti-Sh3Pxd2B Antibody Cat#HPA058141, Table S1) and the omni-tag. The GST-fused proteins were visualized via Coomassie or with antibodies against GST. (**H**) Schematic representation of the TKS4L and TKS4b isoforms and their interaction partners identified in this study. The SH3 domain-binding regions are highlighted in red.

We next examined binding of TKS4L and TKS4b to established TKS4 partners, including the endocytic proteins ITSN1 and ITSN2,^24^ the actin regulator CTTN,^3^ the adaptor GRB2,^25^ SRC kinase,^26^ and TKS4 itself.^23^ For ITSN1 and SRC, both ubiquitously expressed and neuronal SH3 variants (ITSN1 SH3A/SH3An; SRC/nSRC) were tested. Consistent with previous reports, endogenous TKS4L bound the SH3 domains of CTTN, GRB2, and the ubiquitously expressed SRC isoform (Figure 4A), as well as the SH3An, SH3A, SH3C, and SH3E domains of ITSN1 and ITSN2 (Figure 4D,E). Again, no binding of TKS4b to these domains was observed (Figure 4A–E). TKS4L did not interact with the SH3 domains of IRTKS, IRSp53, nSRC, or CSK (Figure 4A), nor with its own SH3 domains, irrespective of the presence (Figure 4F) or absence (Figure 4G) of overexpressed omni-TKS4b.

Together, these data identify six new association partners of the long isoform TKS4L – AMPH1, BIN1, CRKI, NCK1, NCK2, and TKS5 – while indicating that the short isoform TKS4b does not engage these SH3-mediated interactions under the tested conditions. Given that TKS4b lacks the two C-terminal SH3 domains due to alternative 3′ exon usage, the observed bindings are likely dependent on proline-rich motifs within the linker between SH3_3 and SH3_4. These interactions may be critical for invadopodia assembly and function, in line with the central role of TKS4 in invasive structures.^4^

### The TKS4b isoform is phosphorylated at tyrosine residues in MDA-MB-231 cells

The TKS proteins serve as a platform for the recruitment of key players in EGFR signal transduction.^27^ The scaffold protein TKS4L has three known sites (Y25, Y373, and Y508) phosphorylated by SRC tyrosine kinase. Phosphorylation is crucial for its activation and recruitment to invadopodia formation sites.^1^ The TKS4b isoform lacks one of these three tyrosines, but the two remaining residues may potentially be phosphorylated. Therefore, we studied whether the TKS4b isoform is phosphorylated and whether it depends on signaling pathways activated by phorbol ester (PDBu), a potent activator of signaling that promotes invadopodia formation and cell invasion. Lysates of vehicle-or PDBu-stimulated MDA-MB-231 cells overexpressing omni-TKS4b were used for co-immunoprecipitation with antibodies against omni, followed by WB with anti-phosphotyrosine antibodies. Our data demonstrated the phosphorylation of TKS4b in MDA-MB-231 cells only after PDBu treatment and not in unstimulated cells (Figure 5A). Thus, the phosphorylation of TKS4b depends on the activation of signaling pathways associated with invadopodia formation.

**Figure 5.**
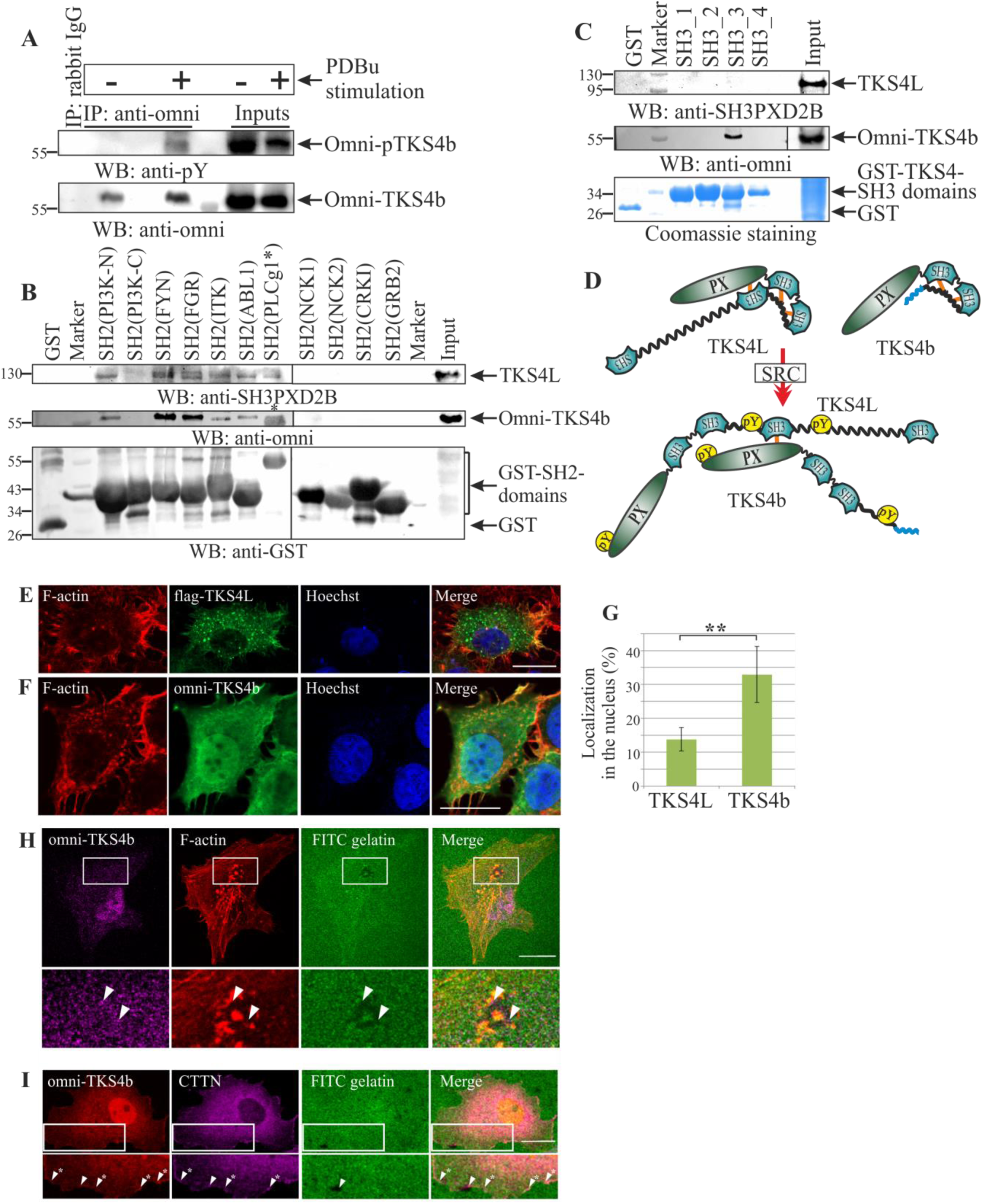
Subcellular localization of the TKS4 isoform and the effect of their phosphorylation on association with binding partners. **(A)** WB analysis of omni-TKS4b phosphorylation in untreated or PDBu-stimulated MDA-MB-231 cell lysates. Rabbit IgG was used as a control. The precipitated proteins were analyzed with anti-phosphotyrosine (pY) and anti-omni antibodies. **(B)** Bacterially expressed GST-SH2 domains of PI3K, FYN, FGR, ITK, ABL, PLCg1, NCK1, NCK2, CRKI, GRB2 and **(C)** GST-SН3 domains of TKS4 were affinity purified and used as bait for pull-down of TKS4 isoforms from the cell extract with tetracycline-induced dominant active myc-Src(Y527F) and transiently expressed omni-TKS4b. Bound proteins were detected by immunoblotting with antibodies against TKS4L or omni-tag. GST-fused proteins were visualized via Coomassie or anti-GST antibodies. * – Nonspecific detection of the GST-SH2 domain of PLCγ1 by anti-omni antibodies, which hinders the identification of the interaction between omni-TKS4b and this SH2 domain. **(D)** Hypothetical interaction of TKS4L and TKS4b isoforms after phosphorylation by SRC kinase. Closed autoinhibited conformation of isoform and interaction of isoforms after phosphorylation. **(E)** MCF-7wt cells were plated on coverslips and transfected with flag-TKS4L or **(F)** omni-TKS4b. TKS4L and TKS4b were detected with anti-flag and anti-omni antibodies, respectively, followed by visualization with an Alexa Fluor 488-conjugated secondary antibody. F-actin was stained with Alexa 555-phalloidin. Nuclei were detected with Hoechst. **(G)** Percentage of TKS4L and TKS4b integrated fluorescence density in nuclei normalized to the total cell area. Two-sample t-test ** – *p* < 0.01. **(H)** The TKS4b protein is localized to sites of invadopodia formation in MDA-MB-231 cells. The TKS4b protein was detected with anti-omni antibodies and visualized with Alexa 633-conjugated secondary antibodies. The arrows indicate the colocalization of TKS4b and F-actin in invadopodia. **(I)** Partial сolocalization of omni-TKS4b and CTTN at the plasma membrane in MDA-MB-231 cells. TKS4b and CTTN were detected with anti-omni and anti-CTTN antibodies and visualized with Alexa 555-conjugated and Alexa 633-conjugated secondary antibodies, respectively. The arrows indicate sites of TKS4b and CTTN colocalization in invadopodia. Arrows with asterisks indicate the colocalization of TKS4b and CTTN in other cellular structures. Scale bar: 20 μm.

Considering the different numbers of tyrosine phosphorylation sites in TKS4L and TKS4b, we sought new binding interfaces that could provide isoform-specific protein-protein interactions. Phosphorylated tyrosine residues in proteins are recognized by the SH2 domains of signaling proteins. The SH2 domains of proteins potentially binding TKS4 were identified via the Scansite service. We also created a T-REx-293 cell line with tetracycline-inducible expression of myc-tagged dominant active mutant Src kinase (Y527F). In this cell line, we additionally transiently expressed omni-TKS4b with or without myc-Src(Y527F) induction. These lysates were used for assays with the GST-fused SH2 domains of the kinases PI3K, FYN, FGR, ITK, and ABL; the adaptor proteins NCK1, NCK2, CRKI, GRB2; and the phospholipase PLCγ1.

We observed that the SH2 domains of PI3K, FYN, FGR, ITK, ABL1, and PLCγ1 efficiently precipitated both TKS4L and TKS4b only from lysates of cells with myc-Src(Y527F) induction (Figure 5B) and not from uninduced cells (Figure S3). No interactions of either the TKS4 isoform with the SH2 domains of the adaptor proteins NCK1, NCK2, CRKI, and GRB2 were detected. Unfortunately, owing to the nonspecific interaction of omni antibodies with the GST-SH2 PLCγ1, it is unclear whether omni-TKS4b interacts with this domain.

We also investigated the possible effects of TKS4L and TKS4b phosphorylation on the interaction with their own SH3 domains and found that phosphorylated omni-TKS4b can interact with the third SH3 domain of TKS4 (Figure 5C), which does not occur in the absence of phosphorylation (Figure 4F-G). Interestingly, the long isoform TKS4L does not acquire this ability after phosphorylation by myc-Src(Y527F).

Although the TKS4b isoform has significantly fewer SH3-containing partners than does TKS4L because of the absence of proline-rich motifs in its structure, the phosphorylation of both TKS4b and TKS4L enables their binding of the SH2 domains of proteins potentially involved in EGFR signaling.

### The TKS4b isoform is localized to the nuclei and invadopodia of human BC cells

The scaffold protein TKS4L is predominantly localized in the cytoplasm. In Src-transformed and invasive cancer cells, it is also found at sites of invadopodia and podosome formation.^1^ The intracellular distribution of the TKS4b isoform is currently unknown. We studied the localization of overexpressed TKS4b in MCF-7 cells and compared it with that of TKS4L (Figure 5E-F). Interestingly, TKS4b was localized in both the cytoplasm and the nucleus. Moreover, its amount in the nuclei of MCF-7 cells was nearly 2.5 times greater than that of the main TKS4L isoform (Figure 5G). Analysis of the amino acid sequence of TKS4b did not reveal nuclear localization signals, suggesting that the mechanisms of its nuclear transport require further investigation.

It has been previously shown that the long isoform TKS4L is a critical component of invadopodia. Its knockdown leads to the formation of incomplete invadopodia.^1^ However, it is unknown whether other TKS4 isoforms participate in this process. As a model for our experiments, we chose MDA-MB-231 BC cells, which are characterized by high invasiveness. We investigated the possible localization of the TKS4b isoform to invadopodia via extracellular matrix proteolytic degradation. During invadopodia formation, the extracellular matrix (FITC-gelatin) is degraded, resulting in black gaps where the fluorescent signal in gelatin is absent. These black gaps, along with F-actin or CTTN staining, were used as invadopodia markers. Immunofluorescence analysis revealed that the omni-TKS4b isoform colocalized with F-actin and CTTN in invadopodia of MDA-MB-231 cells (Figure 5H-I).

Interestingly, the colocalization of TKS4b with CTTN, a marker of active actin cytoskeleton remodeling, was observed not only in invadopodia but also in other cellular structures (Figure 5I).

### Effects of the TKS4L and TKS4b isoforms on the migration and proliferation of MCF-**7 cells**

To evaluate the effects of the TKS4 isoforms on cell proliferation we created MCF-7ΔTKS4, MCF-7+TKS4L, and MCF-7+TKS4b cell lines and performed an MTT assay. Wild-type cells were used as a reference. The MCF-7ΔTKS4 cell line with TKS4 gene knockout was generated via CRISPR/Cas9 technology. The guide sequence for the knockout of the *TKS4* gene was selected in the PX domain-coding sequence of TKS4, which should theoretically stop the synthesis of all known TKS4 isoforms. The cell lines with constitutive overexpression of the long TKS4L isoform (MCF-7+TKS4L) and the TKS4b isoform (MCF-7+TKS4b) were created via the pcDNA4/TO system.

We observed that *TKS4* gene knockout initially led to a significant increase in the cell proliferation rate (1.5-2 times from day 3 to day 5); then, by day 7, the proliferation rate was equal to that of MCF-7wt, and after day 9, it decreased twofold. At the same time, MCF-7+TKS4L and MCF-7+TKS4b cells proliferated at the same rate as the MCF-7wt cells did until day 5 and then slowed 0.5-2 times (Table 1, Figure S4). The results indicate that *TKS4* gene knockout may promote cell proliferation at low cell density, whereas overexpression of both its isoforms does not affect proliferation under the same conditions. It should be noted that MTT assay results become less reliable at high cell density due to contact-mediated growth inhibition. Therefore, the observed decrease in cell proliferation after 5-days in culture cannot be attributed solely to isoforms overexpression.

**Table 1.** The relative optical density (OD^570^) was normalized to that of MCF-7wt (M±σ, n=6)

| Cell line | Relative optical density OD<sup>570</sup>, %, on the day of the experiment |  |  |  |  |
| --- | --- | --- | --- | --- | --- |
|  | 1 | 3 | 5 | 7 | 9 |
| MCF-7ΔTKS4 | 102±39 | 145±8*** | 211±50** | 92±22 | 56±9*** |
| MCF-7+TKS4L | 67±13 | 74±23* | 83±20 | 54±11*** | 49±13*** |
| MCF-7+TKS4b | 103±20 | 75±9** | 112±11 | 87±10* | 75±7*** |

\* – $p < 0.05$ ; \*\* – $p < 0.01$ ; \*\*\* – $p < 0.001$

A number of studies have demonstrated the ability of TKS4 to regulate actin cytoskeleton remodeling and the rate of cell migration.^3,28^ Thus, we investigated the migration ability of these MCF-7 cell lines via the classic scratch test. Indeed, *TKS4* gene knockout reduced the cell migration rate (Figure 6A-B), supporting the idea that *TKS4* is involved in controlling cell motility. In turn, the overexpression of TKS4b led to increased migration, whereas TKS4L overexpression did not affect the migration rate compared with that of wild-type cells.

**Figure 6.**
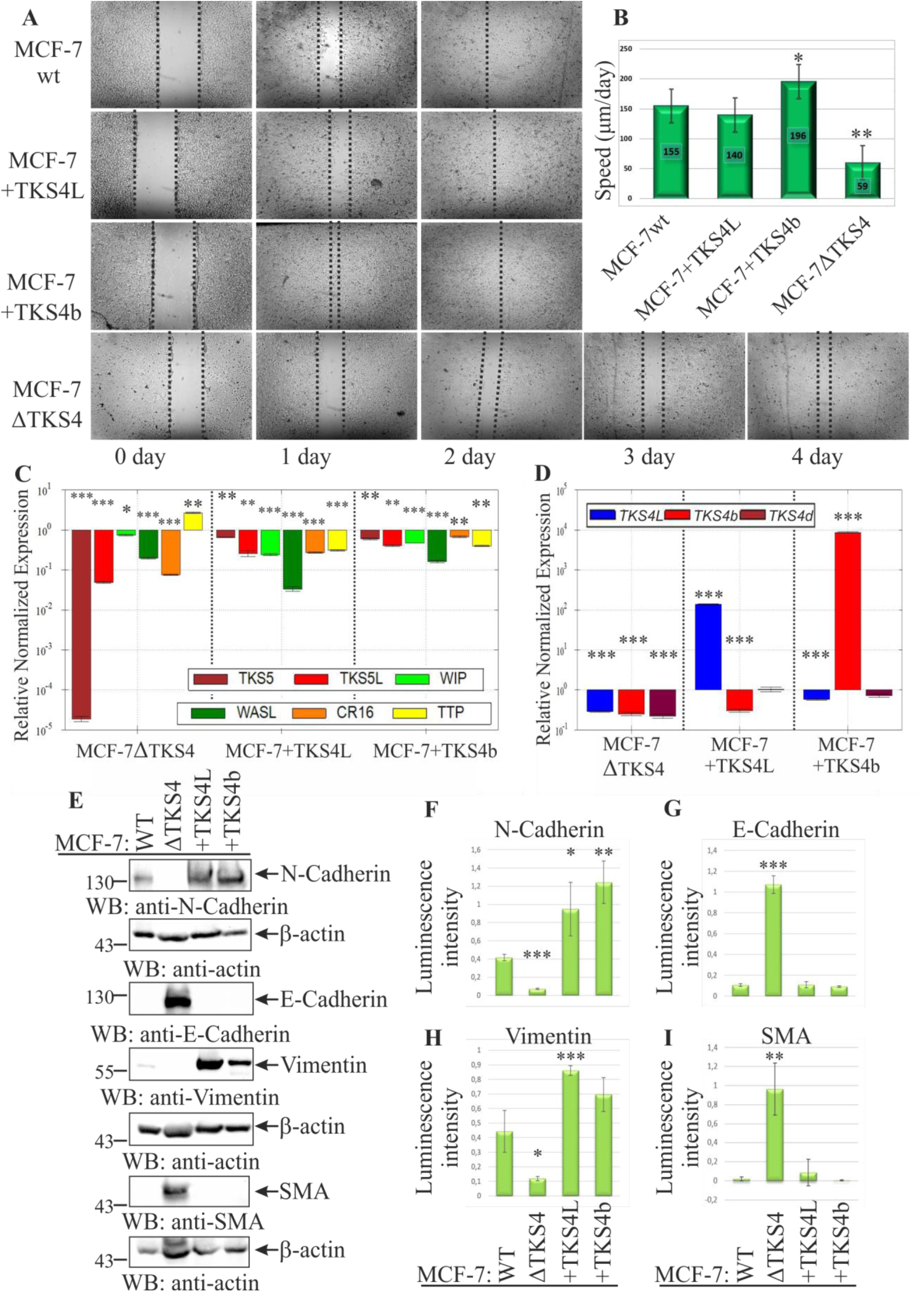
The effect of TKS4 isoforms on the cell migration, epithelial-mesenchymal transition, and the expression of migration-associated genes in MCF-7 cells. **(A)** Cell migration analysis via the scratch test. Confluent MCF-7wt (n=4), MCF-7+TKS4L (n=3), MCF-7+TKS4b (n=3), and MCF-7ΔTKS4 (n=4) cells were scratched, and images were taken immediately after scratching and after 1-4 days. Representative images of wound closure. **(B)** Quantitative comparison of migration rates (μm/day) after 1 day. T-test * – *p* < 0.05, ** – *p* < 0.01 compared with MCF-7wt. Relative expression of genes associated with actin cytoskeleton remodeling (*TKS5*, *WIP*, *WASL*, *CR16*, and *TTP* (**C**)) and (*TKS4L*, *TKS4b*, and *TKS4d* (**D**)) transcripts in the MCF-7ΔTKS4, MCF-7+TKS4L, and MCF-7+TKS4b cell lines compared with the MCF-7wt cell line. The expression ΔΔCq was normalized to that of the *TBP* gene with the standard error of the mean (lg). The data are presented in log10 format. One-way ANOVA * – *p* < 0.05, ** – *p* < 0.01, *** – *p* < 0.001. **(E)** WB analysis of N-cadherin, E-cadherin, vimentin and SMA expression in the MCF-7wt, MCF-7ΔTKS4, MCF-7+TKS4L, and MCF-7+TKS4b cell lines. Graphs of normalized β-actin expression of N-cadherin (**F**), E-cadherin (**G**), vimentin (**H**) and SMA (**I**). Mean±SD (standard deviation), n=3. * – *p* < 0.05; ** – *p* < 0.01; *** – *p* < 0.001 compared with the MCF-7wt.

We subsequently performed qPCR in the stable cell lines MCF-7+TKS4L, MCF-7+TKS4b, and MCF-7ΔTKS4 to assess the expression of genes associated with cell invasion, migration, and actin cytoskeleton remodeling, namely *TKS5*, *WIP*, *WASL*, *CR16*, and *TTP*.^24,27,29,30^ Both the knockout and overexpression of individual TKS4 isoforms exert an inhibitory effect on *TKS5* expression, including the long *TKS5L* transcript as well as the total *TKS5* pool transcripts. *WIP*, *WASL*, and *CR16* expression levels also decreased in all the studied cell lines compared with those in the MCF-7wt. Interestingly, *TKS4* knockout increased the *TTP* expression level, while in MCF-7+TKS4L and MCF-7+TKS4b cells, *TTP* expression was inhibited (Figure 6C, Table S6).

We also investigated the effects of *TKS4L* or *TKS4b* overexpression on the mRNA levels of other currently known *TKS4* transcripts. Compared with those in wild-type cells, *TKS4L* mRNA levels in MCF-7+TKS4b cells were lower. Similarly, in MCF-7+TKS4L cells, *TKS4b* transcript levels decreased. However, the expression of the *TKS4d* transcript did not significantly change in either cell line (Figure 6D, Table S6). Thus, a negative correlation in expression is observed upon the overexpression of *TKS4L* and *TKS4b* isoforms, while the level of *TKS4d* transcripts remains unchanged.

### TKS4L and TKS4b isoforms promote EMT in MCF-7 cells

Previously, *TKS4* gene knockout was shown to lead to EMT in human colon cancer HCT116 cells and lung adenocarcinoma A549 cells.^9,10,31^ EMT is an important process for tumorigenesis and cancer metastasis. We tested whether knockout or overexpression of TKS4 isoforms would induce EMT in BC MCF-7 cells. Initially, luminal A subtype BC MCF-7 cells were derived from the differentiated mammary epithelium origin, but they also presented slight mesenchymal features and increased migration capacity, probably due to malignant transformation.

Epithelial–mesenchymal transition (EMT) is characterized by the loss of epithelial markers, such as E-cadherin and α-smooth muscle actin (SMA), and the acquisition of mesenchymal markers, including N-cadherin and vimentin. We examined the expression of these markers by immunoblotting in MCF-7ΔTKS4, MCF-7+TKS4L, and MCF-7+TKS4b cell lines. Overexpression of either TKS4L or TKS4b was associated with an enhanced mesenchymal phenotype, reflected by increased N-cadherin and vimentin levels. Conversely, MCF-7ΔTKS4 cells displayed elevated E-cadherin and SMA expression, with no detectable N-cadherin or vimentin, indicative of a pronounced epithelial phenotype that exceeded that of parental MCF-7wt cells (Figure 6E–I).

Taken together, these data indicate that TKS4 knockout in MCF-7 cells promotes a mesenchymal–epithelial transition (MET), leading to a more epithelial, less motile phenotype, as further supported by reduced migration in scratch assays. In contrast, overexpression of TKS4L or TKS4b reinforces mesenchymal traits and supports EMT progression.

## DISCUSSION

In this study, we investigated the transcripts of the *TKS4* gene, which encodes a scaffold protein critically important for invadopodia formation. Four *TKS4* mRNA transcripts are formed as a result of alternative splicing, significantly affecting the structure of the proteins. We also compared the expression and functions of its isoforms, which had not been previously studied.

By exploring the levels of *TKS4* transcripts in cell lines, we found that the expression patterns of *TKS4L*, *TKS4b*, and *TKS4d* varied across different cell types (Figure 2, Table S2). However, the expression of individual isoforms did not correlate with the expression of the total *TKS4* mRNA, suggesting the possible existence of yet undiscovered transcripts, which prevents the comparison of the expression levels of known transcripts with the total pool of *TKS4* mRNA.

The TKS4c isoform, generated by the insertion of an additional sequence between the SH3_1 and SH3_2 domains, remains functionally uncharacterized and requires further investigation. Notably, these two domains are known to act cooperatively in binding with the specific TAM proline motif (Figure 1), thereby promoting the formation of a closed autoinhibitory conformation.^23^ It is therefore plausible that elongation of the linker region between them may affect this intramolecular interaction and modulate the autoinhibitory state.

Analysis of *TKS4* or *TKS4L* mRNA expression in different BC tumors revealed that the total pool of *TKS4* transcripts did not differ from that in adjacent tissues, which was correlated with data from the GENT2 and TCGA-BRCA databases. However, total *TKS4* expression levels varied between tumors and were significantly greater in the HER2-Enriched type than in the LumB HER2– and triple-negative types. Moreover, when we analyzed the expression of each isoform separately, we observed that *TKS4L* mRNA was significantly upregulated in all tumor types compared with adjacent tissues. Statistical differences were also demonstrated for the *TKS4L*/*TKS4b* ratio, with an increase in *TKS4L* transcripts in all tumor types compared with adjacent tissues. Thus, despite a constant total *TKS4* expression rate, the *TKS4L* mRNA expression level and/or the ratio of *TKS4L*/*TKS4b* transcripts could be used as a marker for malignancy in human BC tumors. For the same purpose, the expression of individual *TKS4* transcripts, as well as their ratios, should also be investigated in other cancer types.

Before this study, 20 TKS4-interacting proteins were identified via various methods (excluding large-scale mass spectrometry studies, which require further confirmation) (Table 2).

**Table 2.**
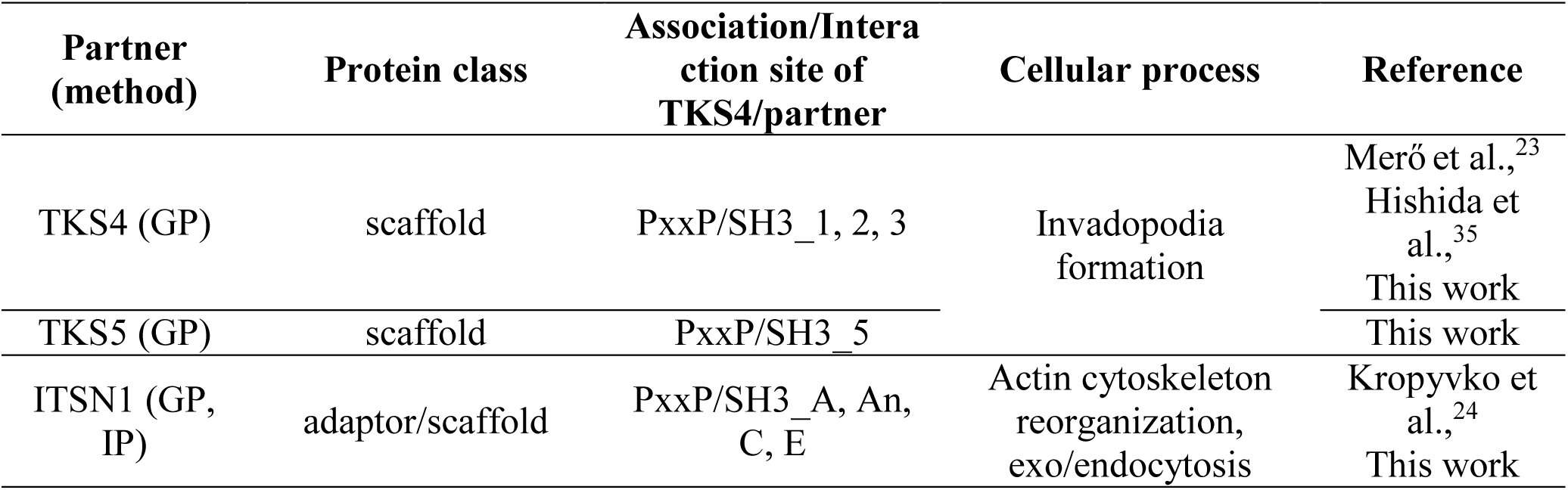
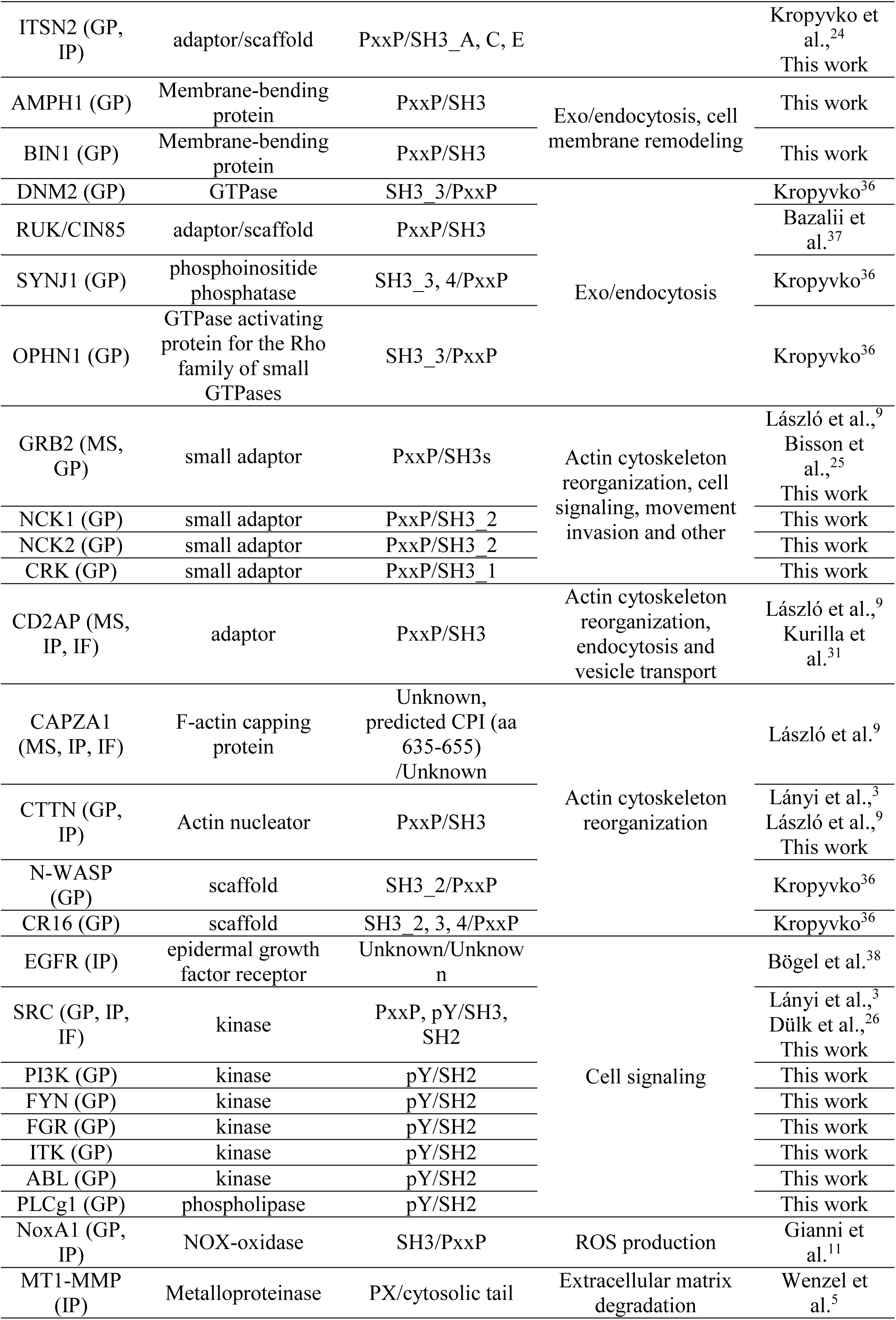
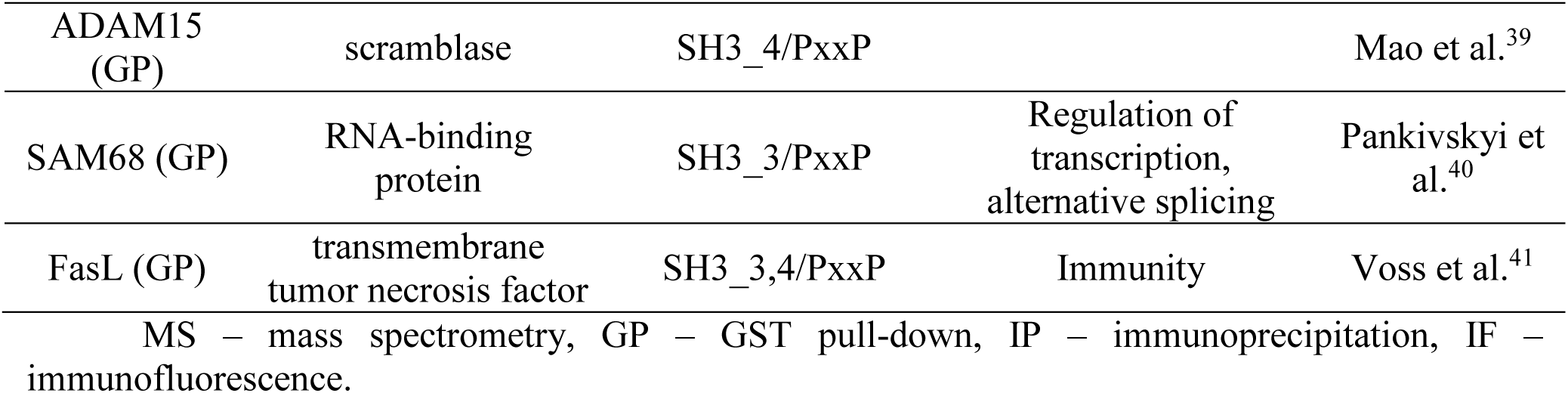
TKS4 associating partners.

In this study, we identified 12 additional protein partners for TKS4L, including another member of the TKS family, TKS5. These findings align with those of Murphy et al., who reported that TKS4L and TKS5 are both involved in common cellular processes, particularly invadopodia formation: TKS5 is involved in the early stages, whereas TKS4 participates in the maturation of invasive structures.^4^

The interaction of TKS4L with the family of membrane-deforming amphiphysin proteins (AMPH1 and BIN1) suggests its role in endocytosis, exocytosis, and vesicular transport. Interactions with the small adaptors NCK1, NCK2, and CRK link TKS4L to cell movement, signal transduction, and other processes involving these proteins. This may occur in conjunction with TKS5, which also interacts with NCK1 and NCK2 and regulates migration and invasion through the Src-Tks5-Nck pathway.^42^

Interestingly, the newly identified SH3-containing partners of TKS4 associate exclusively with the long isoform TKS4L and not with TKS4b, highlighting the importance of proline-rich motifs located in the linker region between the 3rd and 4th SH3 domains, which are absent in the TKS4b isoform (Figure 4H). Here, we did not study the reverse associations – those of protein partners with proline-rich motifs associating with the SH3 domains of TKS4 – but most of the previously described TKS4 partners also associate predominantly with its 3rd and/or 4th SH3 domains (Table 2), making interaction with TKS4b impossible. Among the potential SH3-binding partners for TKS4b, we envision the actin-regulatory proteins N-WASP and CR16. These proteins interact with the second SH3 domain of TKS4L, which is also present in TKS4b.^36^

The TKS4L isoform is phosphorylated in the 3T3 cell line at three sites, Y25, Y373, and Y508, which are important for its activation and involvement in invadopodia formation.^1^ TKS4b has two of the three tyrosines, but its phosphorylation has not yet been studied. We observed a high level of TKS4b phosphorylation after stimulation with PDBu, a known inducer of invadopodia formation. The TKS4L protein is a substrate for SRC kinase, and their interaction is mediated by proline motifs (aa 466-474) together with the phosphorylated Y508 of TKS4 and the SH3 and SH2 domains of SRC kinase.^26^ Interestingly, TKS4L interacts with the neuron-specific SH3A domain of ITSN1 and does not bind the neuron-specific isoform SH3 (nSRC) of the SRC kinase. This may indicate other mechanisms of TKS4 phosphorylation in neurons. The TKS4b isoform cannot form a complex with SRC kinase in the absence of binding sites, but phosphorylation by this kinase still occurs.

TKS4 phosphorylation leads to the transition of the TKS4 molecule from a closed inactive state to an activated open state. Mero et al. demonstrated that TKS4L adopts a closed autoinhibitory conformation through interactions of the first two SH3 domains and the TAM located before the third SH3 domain, with the SH3_3 domain interacting with the PX domain.^23^ This typical domain organization and autoinhibition can be observed in all five members of the p47phox-related organizer superfamily.^23^ We believe that the regulation of TKS4b activity may also occur through intramolecular interactions, as most of the necessary sites are present in its structure (Figure 1). We observed that the unphosphorylated TKS4b isoform does not bind to the SH3_3 domain of TKS4 (Figure 4F), but the interaction is established after phosphorylation by SRC kinase (Figure 5C). This is likely due to the transition of TKS4b to the open activated conformation upon phosphorylation at Y373, which is located immediately after TAM, and the release of the interaction site in the PX domain (Figure 5D).

Proteins that compete for interaction with the SH3_1 and SH3_2 domains of TKS4, such as N-WASP, may also act as activators of TKS4. However, the lack of interaction between phosphorylated TKS4L isoform and its SH3_3 domain cannot currently be explained. We did not observe another expected interaction between the phosphorylated TKS4L or TKS4b and SH3_1 and SH3_2 domains. The open conformation should theoretically allow for dimer formation between these domains and TAM from another TKS4L or TKS4b molecule. The absence of such interactions in our experiments may be explained by the fact that, as shown by Mero et al., both SH3_1 and SH3_2 domains are simultaneously required for TAM binding, whereas we used isolated domains.^23^ Thus, we suppose that the formation of mono-or heterodimers of open conformation TKS4L and TKS4b isoforms is still possible, but this issue needs further confirmation (Figure S5).

Additionally, the phosphorylation of TKS4b and TKS4L by SRC kinase promotes their interaction with the SH2 domains of proteins involved in EGFR signaling, namely, the kinases PI3K, FYN, FGR, ITK, and ABL1 and the phospholipase PLCγ1. This finding suggests a broader role for TKS4 in cellular signaling networks, with both the TKS4L and TKS4b isoforms likely contributing to these processes.

The regulation of TKS4 binding to PtdInsP could also be regulated by the phospholipase PLCγ1, which binds to phosphorylated TKS4 through its SH2 domain (Figure 5B). PLCγ1 cleaves phosphatidylinositol bisphosphates into inositol-1,4,5-triphosphate and diacylglycerol, which may not only regulate the interaction of the PX domain of TKS4 isoforms with the membrane but also recruit them to signaling pathways regulating proliferation, migration, apoptosis, and other cellular processes.^43,44^

Next, we investigated the functional features of the TKS4b and TKS4L isoforms. Compared with wild-type cells, overexpression of TKS4b in the MCF-7 cell line resulted in increased cell migration but had no effect on proliferation. Interestingly, compared with cells overexpressing TKS4L, MCF-7+TKS4b cells exhibited increased cell migration. Knockout of the *TKS4* gene in MCF-7ΔTKS4 cells resulted in increased proliferation and decreased migration compared with wild-type cells. The findings suggest at least partial involvement of TKS4 isoforms in the regulation of cell movement and proliferation. Therefore, we studied the underlying cellular processes, such as the expression of genes associated with cell migration and invasion. Surprisingly, both knockout and overexpression of individual TKS4 isoforms significantly reduced *TKS5*, *TKS5L*, *WIP*, *WASL*, *CR16*, *TKS4b*, and *TKS4L* levels. However, the expression of the RNA-binding protein *TTP* gene was upregulated upon *TKS4* gene knockout but downregulated upon overexpression of TKS4 isoforms. According to previous studies, TTP overexpression decreases cell motility and invasion in the MDA-MB-231 cell line, which is consistent with our data.^17^ Thus, we suppose that the reduction in cell migration, which we observed in MCF-7 cells upon *TKS4* gene knockout, could be linked to an increase in *TTP* levels.

Notably, MCF-7 is the least aggressive noninvasive human BC cell line with the most common LumA molecular subtype.^45^ In contrast, MDA-MB-231 cells are invasive cells that correspond to the triple-negative BC subtype. As we demonstrated, the expression levels and ratios of *TKS4L* and *TKS4b* transcripts differ even in these closely related cells, suggesting that the TKS4L and TKS4b isoforms may play distinct roles in cellular function.

The TKS4L scaffold is one of the key proteins in invadopodia.^4^ It takes part in actin polymerization and membrane bending processes. Together with the actin nucleator CTTN, TKS4L plays a central role in the maturation of invadopodia by regulating the secretion and recruitment of membrane metalloproteinases to sites of invasive structure formation, which leads to extracellular matrix degradation and enables cell migration and invasion into surrounding tissues.^46,47^ Here, we demonstrated the localization of the TKS4b isoform in the invadopodia of MDA-MB-231 cells together with CTTN, indicating the involvement of TKS4b in the development of invasive structures. By performing functional investigations, we observed different subcellular distributions of TKS4b and TKS4L. TKS4b significantly accumulated in the nucleus, whereas TKS4L was mainly present in the cytosol of both the MCF-7 and MDA-MB-231 (the results are shown only for the MCF-7) model cell lines. This finding is intriguing, as there are no known nuclear localization signals in the TKS4b sequence. Its function in the nucleus also remains unclear.

The EMT, during which epithelial cells transform into a mesenchymal state, is a central regulator of metastasis.^48^ Given the implication of TKS4 isoforms in cell motility and invasion, we hypothesized that it could also be involved in EMT, as it is closely linked to both of these processes.

Analysis of EMT marker expression in cells with TKS4 isoform overexpression and knockout revealed that overexpression of both the TKS4L and TKS4b isoforms enhanced the mesenchymal properties of MCF-7 cells, whereas *TKS4* gene knockout resulted in an epithelial-like phenotype. These results are consistent with our data showing reduced motility of MCF-7ΔTKS4 cells. However, our findings differ from those from studies conducted on A549 lung cancer cells and HCT116 colon cancer cells, where *TKS4* gene knockout enhanced mesenchymal characteristics, increasing the expression of mesenchymal markers and cell motility.^9,10,28,31,49^ We hypothesize that *TKS4* gene dysfunction in various cell lines could lead to different functional outcomes. Therefore, the role of TKS4 isoforms in one cell type cannot be generalized to others, and each cell model must be assessed individually via isoform-specific nucleotide sequences for knockout or siRNA experiments. Different expression levels and ratios in various cell lines (Figure 2) may indicate the independent role of these isoforms in both normal cellular physiology and in diseases associated with *TKS4* gene dysfunction. A limitation of this study is that all functional experiments were performed exclusively in the MCF-7 BC cell line. Although MCF-7 represents a well-established luminal A subtype and exhibits the highest expression of *TKS4L* and *TKS4b* among the cell lines analyzed, future studies should investigate the roles of these isoforms in BC cell lines with different molecular characteristics and invasive capacities to determine whether their functions are conserved across BC subtypes.

Our data contribute to the understanding of the expression and functions of the TKS4L and TKS4b scaffold isoforms in BC and raise new issues in considering the role of alternative splicing and differential protein isoform levels in various cellular models.

### Limitations of the study

One limitation of this study is the relatively small sample size of the breast cancer cohorts analyzed for *TKS4* transcripts expression. Larger sample sizes may reveal additional statistically significant differences in isoform expression, particularly for *TKS4b*.

Breast cancer is a heterogeneous disease comprising multiple molecular subtypes. Therefore, further studies using additional breast cancer cell lines, including triple-negative and HER2+ models, will be necessary to determine whether the regulatory relationships between *TKS4* transcript isoforms observed in MCF-7 cells are conserved across different breast cancer contexts.

Another limitation is the absence of *in vivo* validation of the identified protein complexes involving TKS4 associated partners. Such studies would be essential to confirm these complexes and to further characterize their biological relevance.

## RESOURCE AVAILABILITY

### Lead contact

Further information and requests for resources and reagents should be directed to and will be fulfilled by the lead contact, Serhii Kropyvko.

### Materials availability

All materials from this study will be provided upon request to the corresponding author, Serhii Kropyvko.

### Data availability

The datasets used or analyzed in the present study are available from the corresponding author upon reasonable request. The results of the qPCR analysis of gene expression data are available from the corresponding author upon reasonable request or have been uploaded to the GEO database GSE304284 (https://www.ncbi.nlm.nih.gov/geo/query/acc.cgi?acc=GSE304284).

## ACKNOWLEDGMENTS

We are extremely grateful for our success, which would not have been possible without the support and care of Prof. Alla Ryndich (Kyiv, Ukraine). We are grateful to Dr. Oleksandr Dergai for the critical reading of the manuscript. We are grateful to Prof. Sara A. Courtneidge (La Jolla, USA), Dr. Mykola Dergai (Kyiv, Ukraine) and Dr. Oleksandr Dergai (Kyiv, Ukraine) for providing genetic constructs. We are grateful to Dr. Serhii Karakhim (Kyiv, Ukraine) for providing expertise and help in obtaining the confocal images. We would like to express our gratitude to the Ukrainian Armed Forces, friends, and partners of Ukraine who made this work possible.

This research was partially supported by a grant from the Simons Foundation (Award #1290589, [Serhii Kropyvko and Tetyana Gryaznova]) and the National Research Foundation of Ukraine Grant (No. 2020.01/0021).

## AUTHOR CONTRIBUTION

**S.K.**: Project administration; Data curation; Conceptualization; Formal analysis; Investigation; Validation; Writing-original draft. **N.S.**: Formal analysis; Investigation; Software. **O.G.**: Formal analysis; Investigation; Validation; Writing-original draft. **K.L.**: Investigation; Visualization. **Y.N.**: Investigation; Visualization. **D.K.**: Investigation; Resources. **V.P.**: Investigation; Visualization. **V.K.**: Investigation; Visualization. **L.S.**: Investigation; Resources. **N.V.**: Investigation; Resources. **T.G.**: Investigation; Methodology; Validation; Writing-original draft.

## SUPPLEMENTAL INFORMATION

Figures S1-S6 and Tables S1-9.

## METHODS

### Cell culture and transfection

The following human cell lines were used in this study: embryonic kidney 293; renal cell carcinoma KRC/Y; cervical cancer HeLa; retinal ARPE-19; breast cancer T-47D, MCF-7 and MDA-MB-231; glioblastoma U-251 MG, U-87 MG and T98G; neuroblastoma SH-SY5Y; ovarian adenocarcinoma SKOV-3; fibrosarcoma HT1080; normal skin fibroblasts HaCat; skin epidermoid carcinoma A431; cutaneous melanoma WM793; hepatocellular carcinoma Hep G2; colorectal adenocarcinoma HT-29; lung carcinoma A549; testicular seminoma JKT-1; normal prostate PNT2; prostate cancer DU145, LNCaP and PC-3; urinary bladder carcinoma 5637 and T24 and Flp-In™ T-REx™-293. For more information, see Supplementary Table S7.

Cell lines were authenticated for the described experiments and have not been previously reported as misidentified or contaminated. Cell lines were free of mycoplasma contamination for the described experiments.

The cells were cultured in Dulbecco’s modified Eagle’s medium (DMEM) supplemented with 10% fetal bovine serum (Sigma-Aldrich, F7524-500ML), 50 U/ml penicillin and 100 mg/ml streptomycin at 37°С, 5% CO_2_ in a humidified incubator. The cells were transiently transfected with PEI (polyethyleneimine) or JetPEI reagents and processed 24-48 h after transfection.

MDA-MB-231 cells were transfected and 24 h after serum starved for 16 h and then restimulated with PDBu (100 μg/ml) for 30 min.

### Generation of MCF-7 stable cell lines with a knockout and overexpression of the *TKS4* gene

MCF-7 cells with a knockout of *TKS4* gene were obtained CRISPR/Cas9-mediated gene editing.^50^ The sequences of gRNA oligonucleotides – 5’gatgttggacaaatttccca3’. Transfection of pSpCas9(BB)-2A-Puro(PX459)V2.0-TKS4 into MCF-7 cells was carried out using jetPEI transfection reagent. The cell culture dishes 24 hours posttransfection selected with puromycin (3 μg/ml) for 48-72 hours.

Stable cell lines with overexpression of the TKS4L isoform (MCF-7+TKS4L) and TKS4b isoform (MCF-7+TKS4b) were generated via the pcDNA4/TO system according to the manufacturer’s instructions. MCF-7 cells were transfected with the pcDNA4/TO-myc-TKS4L and pcDNA4/TO-omni-TKS4b plasmids for 24 h. Afterward, cell selection was carried out for 48-72 h by adding the antibiotic zeocin (300 μg/ml).

Single-cell clones were isolated and expanded and then verified by WB analysis for *TKS4* gene knockout or overexpression of myc-TKS4L and omni-TKS4b isoforms, respectively (Figure S6).

### Generation of doxycycline-inducible Flp-In T-REx-HEK-293 cell lines

To obtain Flp-In T-REx-HEK-293 cells overexpressing omni-TKS4b or myc-Src(Y527F), the pcDNA5/FRT/TO-omni-TKS4b or pcDNA5/FRT/TO-myc-Src(Y527F) vectors were co-transfected together with the pOG44 expression vector into cells using PEI. Cells were selected by adding 150 μg/ml hygromycin for 48 hours, and this process was repeated 2-3 times. Recombinant protein synthesis was induced by 0.5-2 μg/ml doxycycline in the medium for 24 hours.

### Expression constructs

The construct encoding the sequence of the *TKS4* 3rd exon 5′gatgttggacaaatttccca3′ was subcloned into the pSpCas9(BB)-2A-Puro(PX459)V2.0 vector as described in Ran et al.^50^ The cDNA fragments encoding human wild-type full-length TKS4b (residues 2-430, GenBank Accession NO. NP_001295104.1) were subsequently subcloned into the pcDNA4/HisMaxB, pcDNA4/TO and pcDNA5/FRT/TO vectors with an additional omni-tag at the 5′-end. The cDNA fragments encoding human wild-type full-length TKS4L (residues 2-911, Accession NO. NP_001017995) was subcloned into the pcDNA4/TO vector with an additional myc-tag at the 5′-ends, respectively. The cDNA fragment encoding the dominantly active mutant of Src(Y527F) kinase was subcloned into the pcDNA5/FRT/TO vector with an additional myc-tag at the 5′-end. The sequences encoding the SH2 domains of human NCK1 (residues 274-353, Accession NO. NP_006144.1) and NCK2 (residues 281-360, Accession NO. NP_003572.2), the full-length human GRB2 (residues 2-217, Accession NO. NP_002077), the 1st and 2nd SH3 domains of human NCK1 (residues 2-70 and 104-166), the 1st, 2nd and 3rd SH3 domain of human NCK2 (residues 2-66, 108-176 and 195-261) and the 1st, 2nd, 3rd and 5th SH3 domains of human TKS5 (residues 165-226, 149-209, 419-481 and 1043-1104, Accession NO. NP_055446.2) were subcloned into the pGEX-4T-1 vector (Amersham Biosciences, 27-4580-01). The sequences encoding the 3rd SH3 domains of human NCK1 (residues 188-255), the SH3 domain of human IRSp53 (residues 371-438, Accession NO. NP_059345.1) and the 4th SH3 domain of human TKS5 (residues 815-873) were subcloned into the pGEX-4T-2 vector. Sequences encoding the SH3 domains of human IRTKS (residues 339-402, Accession NO. NP_061330.2), CTTN (residues 479-550, Accession NO. NP_005222.2) and BIN1 (residues aa 516-593, Accession NO. NP_647593.1) were subcloned into the pGEX-4T-3 vector. The pOG44 vector was used for cotransfection with the pcDNA5/FRT/TO vector. Flag-TKS4L was a kind gift from Prof. Sara A. Courtneidge (La Jolla, USA). Plasmids encoding the GST-fused SH3 domains of human AMPH1 and neuronal nSRC were a kind gift from Dr. M. Dergai (Kyiv, Ukraine), and the GST-fused SH3 domains of human SRC, CRK, PLCg1 and CSK were a kind gift from Dr. O. Dergai (Kyiv, Ukraine). Constructs encoding the GST-fused SH3 domains of human ITSN1, ITSN2,^51^ TKS4,^36^ and the GST-fused SH2 domains of human GRB2, CRK, ITK, FGR, FYN, ABL1, PI3K1-N, PI3K1-C and PLCg1^52^ were described previously.

### Antibodies and immunofluorescence reagents

Primary antibodies: anti-omni (D-8), 1:5000; anti-Vimentin (V9), 1:1000; anti*-*SMA (CGA7), 1:1000; anti-myc (9E10), 1:5000; anti-phosphotyrosine (PY 99), 1:1000; anti-omni (M-21), 1:5000; anti-flag clone M2 (F1804), 1:5000; anti-CTTN (p80/85) clone 4F11, 1:3000; anti-β-actin clone AC-15, 1:5000; anti-α-tubulin clone B-5-1-2, 1:5000; anti-SH3PXD2B, 1:3000; anti-GST, 1:3000; N-Cadherin, 1:1000 and E-Cadherin, 1:1000. Secondary horseradish peroxidase-labeled antibodies were anti-goat (1:10000), anti-rabbit (1:10000), and anti-mouse (1:10000). For immunofluorescence, goat anti-mouse Alexa Fluor 488 (1:400), goat anti-mouse Alexa Fluor 633 (1:400), donkey anti-rabbit Alexa Fluor 555 (1:400) antibodies and phalloidin Alexa Fluor 555 (1:400) were used. For more information, see Supplementary Table S8.

### Total RNA isolation from human breast tumor samples and cell lines

The study protocol was approved by the Bioethics Commission of the Institute of Molecular Biology and Genetics, NAS of Ukraine (Protocol No. 47, dated 15 July 2025). All patients and healthy donors provided written informed consent before enrollment and were treated in accordance with the ethical and legal principles of the Declaration of Helsinki.

Breast tumor samples were obtained from the National Cancer Institute (Kyiv, Ukraine), immediately frozen in liquid nitrogen after surgical excision, and stored at −80°C until further use. All participants provided written informed consent before sample collection.

Total RNA was isolated from 0.2-1.5 g of tissue or 5×10^6^ human cells via the guanidinium isothiocyanate method via the innuSOLV, TRIzol™ or RNA Go reagents in accordance with the manufacturer’s recommendations.

### cDNA synthesis and qPCR with fluorescence labeled probes

Two to eight micrograms of total RNA were pretreated with DNase I according to the manufacturer’s recommendations to remove residues of genomic DNA. After that, 20 μl of cDNA was synthesized via RevertAid H Minus Reverse Transcriptase according to the manufacturer’s recommendations. The cDNAs were stored at −20°C. qPCR was performed in a 25 μl mixture containing 0.2 μM of each specific primer and 0.1 μM Taq-Man probe, 1.5 mM MgCl_2_, 0.2 mM dNTPs, 2.5 units of DreamTaq DNA polymerase and the corresponding buffer. Amplification was performed under the following conditions: denaturation – +95°С, 20 sec (2 min for the first cycle); primers reassociation and synthesis were combined at +60°C for 1 min, for 50 cycles. Each sample was analyzed in duplicate or triplicate. qPCR was performed with a CFX96 amplifier (Bio-Rad). The *TBP* gene was selected as a reference gene based on the analysis of literature sources that demonstrated its suitability as a control for gene expression analysis in BC.^53,54,55^ For the primers and Taq-Man probes used for qPCR, see Supplementary Table S9.

### Protein expression and purification, GST pull-down assay and Western blot analysis

The recombinant GST-fused proteins were expressed in *Escherichia coli TOP10* and affinity purified via glutathione-Sepharose 4B according to the manufacturer’s instructions. GST-tagged proteins (5-20 μg) coupled to beads or GST alone were incubated for 2 h at 4°C with lysates of human cell lines. Cell lysates were prepared in pull-down extraction buffer containing 20 mM Tris-HCl pH 7.5, 1% Triton, 150 mM NaCl, 10% glycerol, 1 mM EDTA and protease inhibitor cocktail and further cleared by centrifugation for 20 min at 14000 rpm at 4°C. To work with the phosphorylated proteins, we used IP extraction buffer containing 20 mM Tris-HCl pH 7.5, 0.5% NP40, 150 mM NaCl, 10% glycerol, 1 mM Na_3_VO_4_ and protease inhibitor cocktail. The beads with protein complexes were washed with extraction buffer without inhibitors 3-4 times at 4°C and boiled in Laemmli buffer (150 mM Tris-HCl pH 6.8, 2.5% glycerol, 10% SDS, 3% β-mercaptoethanol and 0.5% bromophenol blue). The proteins were separated by SDS-PAGE, transferred to nitrocellulose membranes and blocked in 5% nonfat milk. The membranes were incubated with appropriate primary antibodies, followed by incubation with peroxidase-conjugated secondary antibodies. Immunoreactive bands were detected via enhanced chemiluminescence (ECL) reagents. Chemiluminescence was captured with Molecular Imager ChemiDoc TM XRS+ (Bio-Rad). The signal intensities were quantified via ImageLab^TM^ software. Date visualization and statistical analysis were performed in MS Excel or OriginPro 9.1.

### Co-immunoprecipitation

For co-immunoprecipitation, the cells were lysed in IP extraction buffer (20 mM Tris-HCl pH 7.4, 150 mM NaCl, 1% NP40, 10% glycerol, 2 mM EDTA) with protease inhibitor cocktail. The lysates were incubated with antibodies and protein A/G PLUS-Agarose. After incubation for 3 h at 4°C, the beads with protein complexes were washed 3-4 times with IP buffer without protease inhibitors. Bound proteins were eluted by boiling in Laemmli sample buffer and then analyzed by SDS-PAGE and Western blot.

### Immunofluorescence and confocal microscopy

The cells were plated on coverslips and transfected with the appropriate plasmid DNA via the JetPEI reagent. Twenty-four hours posttransfection, the cells were washed with PBS, fixed in 4% formaldehyde for 15 min and washed three times in PBS containing 0.2% Triton X-100. Non-specific binding was blocked by incubation with 2% BSA and 0.2% Triton X-100 in PBS for 30 min at room temperature followed by incubation with the appropriate antibodies. Cell nuclei were stained with Hoechst. The samples were Mowiol medium containing 2,5% DABCO (Sigma). Confocal images were taken via Zeiss LSM510 confocal microscope with Zeiss LGM Image Browser software (Version 4.0.0.241; Carl Zeiss AG).

### MTT-test

The MCF-7wt, MCF-7ΔTKS4, MCF-7+TKS4L and MCF-7+TKS4b cells were seeded at 1×10^3^ cells per well in 96-well plates and cultured in DMEM supplemented with 10% fetal bovine serum. Each cell line was divided into 5 groups, each containing six technical replicates. The first group was cultured for 1 day and defined as the starting point (0 days). The cells in the other groups were cultured for 3, 5, 7, or 9 days. Afterward, 10 μL of a 5 mg/ml solution of 3-(4.5-dimethylthiazol-2-yl)-2.5-diphenyl tetrazolium bromide (MTT) was added to the cells, followed by incubation for 3 h at +37°C. The formed formazan crystals in the wells were dissolved in 100 μL of DMSO. The optical density was measured at 570 nm using a BioTek ELx800 (BioTek Instruments). To calculate the relative level of cell proliferation, the values obtained for the points corresponding to each cultivation period were divided by the optical density obtained for the initial point (0 days), whose relative absorbance value was set as 1. Statistical analysis was performed via Student’s t-test. All calculations were performed in “RStudio 4.3.1” via the “rstatix” and “tidyverse” packages.

### Scratch test

MCF-7wt, MCF-7ΔTKS4, MCF-7+TKS4L and MCF-7+TKS4b cells were seeded in 6-well plates and grown until 100% confluence to form a monolayer. A scratch was created with a pipette tip across the diameter of the dish. The cells were washed once with PBS and then replaced with fresh medium. At time points of 0, 24, 48 and 72 h of incubation, the cells were examined using Leica DM 1000 microscope. The images were acquired and further analysed quantitatively by ImageJ that measured the distance (µm) travelled by the cells over time. Each experiment was repeated minimum three times.

### Invadopodia gelatin degradation assay

Coverslips were acid-washed and coated with poly-L lysine for 20 min, washed with PBS (phosphate-buffered saline) and crosslinked with 0.5% glutaraldehyde for 15 min. The coverslips were then inverted onto Oregon Green 488-conjugated fluorescent gelatin for 15 min in the dark. After being washed with PBS, the coverslips were incubated with 5 mg/ml sodium borohydride for 3 min at room temperature, followed by another wash with PBS and sterilization with 70% ethanol for 15 min. Finally, the fluorescent gelatin-coated coverslips were incubated with complete growth medium for 1 h at 37°C before being plated. MDA-MB-231 cells were cultured on fluorescent gelatin-coated coverslips for 6 h and then fixed with 4% formaldehyde, blocked with 2% bovine serum albumin (BSA) in PBS containing 0.1% Triton X-100, incubated with the appropriate primary antibodies, washed, incubated with the appropriate fluorescent secondary antibodies and/or phalloidin (F-actin staining), and mounted with Mowiol. The invadopodia were identified by gelatin degradation sites that coincided with F-actin/CTTN staining.

### Bioinformatics analysis

The nucleotide sequences encoding *TKS4* were analyzed via the services of the National Center for Biotechnology Information (NCBI)^56^ and the Ensembl Genome Browser.^57^ For pancancer analysis, we utilized expression data from the Gene Expression Database of Normal and Tumor Tissues 2 (GENT2) database for the *TKS4* gene across multiple tumor types to visualize varying expression levels. The results from the GENT2 database, were used to generate Supplementary Table S3, which ranks gene expression on the basis of p-values and log2-fold change (log2FC) values.^58^ We obtained RNA sequencing (RNA-seq) expression data of the *TKS4* gene for different human breast cancer types (PAM50_mRNA_nature2012 for Fig.3B and PAM50Call_mRNAseq for Fig.3C) from The Cancer Genome Atlas (TCGA) via the University of California Santa Cruz (UCSC) Xena and the SpliceSeq bioinformatic tool respectively.^59,60,61,62^

### QUANTIFICATION AND STATISTICAL ANALYSIS

Analysis of TKS4L and TKS4b nuclear localization was performed with ImageJ software via confocal images of the middle plane of the cells (n=14 for TKS4L and n=17 for TKS4b). The integrated fluorescence density of TKS4L or TKS4b was calculated in the nuclei (the nuclei area was delineated with Hoechst staining) and compared to the fluorescence density in the total cell area (delineated with F-actin staining). The data are presented as the mean percentage of nuclei with standard deviation. Statistical analysis was performed with a two-sample t-test.

All results are expressed as the mean ± standard deviation (S.D.) from three independent experiments. Statistical significance between two groups was determined by using Student’s *t*-test. The difference between groups was considered to be statistically significant at *p* < 0.05, *p* < 0.01 and *p* < 0.001.

### Quantitative PCR calculation

For the qPCR analysis, a CFX96 real-time PCR amplifier (Bio-Rad) with CFX Maestro 2.3 Version 5.3.022.1030 software was used. The following formula was used to calculate the qPCR results: Exp = 2^(E_ref_ ^-Ct(ref)^-E_target_^-Ct(target)^), where E_target_ is the PCR efficacy for the target gene, E_ref_ is the PCR efficacy for the reference gene, ^Ct(target)^ is the mean cycle value for the target gene, and ^Ct(ref)^ is the mean cycle value for the reference gene. The PCR efficacy was determined via LinRegPCR Version 2021.2 software. The cycle values for the target and reference genes were determined via a regression approach built into the software. Statistical processing of the qPCR data was carried out with CFX Maestro 2.3 Version 5.3.022.1030 software via the One-way ANOVA, Fisher’s criterion, and Tukey’s HSD test.

## Supplementary Materials

**Table S1.** Commercial antibodies to TKS4, related to Introduction

| Name | Firm | Catalog number | Host species | Monoclonal/<br>polyclonal | Immunogen<br>region/epitope |
| --- | --- | --- | --- | --- | --- |
| Anti-Sh3Pxd2B Antibody<br>Produced In Rabbit * | Sigma Aldrich | Cat#HPA058141 | Rabbit | Polyclonal | aa 431-532 |
| SH3PXD2B Polyclonal<br>Antibody | Invitrogen | #PA5-57673 | Rabbit | Polyclonal | aa 556-642<br>(#RP-96944) |
| SH3PXD2B Antibody | Novus<br>Biologicals | NBP3-25137 | Rabbit | Polyclonal | aa 431-532 |
| Anti-SH3PXD2B antibody | Abcam | ab122342 | Rabbit | Polyclonal | aa 550-650 |
| Anti-SH3PXD2B | Atlas<br>Antibodies | HPA036471 | Rabbit | Polyclonal | aa 556-642 |
| SH3PXD2B polyclonal<br>antibody | Abnova<br>Corporation | PAB23008 | Rabbit | Polyclonal | aa 556-642 |
| SH3PXD2B antibody (AA<br>300-350) | Antibodies<br>online | ABIN7451528 | Rabbit | Polyclonal | aa 300-350 |
| SH3PXD2B antibody (AA<br>505-539) | Antibodies<br>online | ABIN6244203 | Rabbit | Polyclonal | aa 505-539 |
| SH3PXD2B antibody (AA<br>596-628) | Antibodies<br>online | ABIN6244202 | Rabbit | Polyclonal | aa 596-628 |
| Polyclonal Rabbit<br>anti-Human SH3PXD2B<br>Antibody (aa300-350) | LS Bio | LS-C290107 | Rabbit | Polyclonal | aa 300-350 |
| anti-SH3PXD2B antibody | MyBioSource | MBS6499363 | Rabbit | Polyclonal | aa 257-477 |

\* - antibodies used in this work

**Table S2.** Ratios of *TKS4* gene transcript expression in human cell lines, related to Figures 2.

| **Human cell lines** | ***TKS4L/TKS4b*** | ***TKS4L/TKS4d*** | ***TKS4b/TKS4d*** |
| --- | --- | --- | --- |
| 293 | 0.716 | 3.201 | 4.47 |
| KRC/Y | 0.42 | 0.354 | 0.843 |
| HeLa | — | — | — |
| U-251 MG | 2.031 | 8.835 | 4.348 |
| U-87 MG | 0.923 | 2.05 | 2.221 |
| T98G | 0.939 | 4.357 | 4.637 |
| PNT2 | 1.54 | 4.543 | 2.949 |
| LNCaP | 1.988 | 4.196 | 2.11 |
| DU145 | 0.671 | 0.671 | 1 |
| PC-3 | 1.074 | 6.724 | 5.015 |
| SH-SY5Y | 2.544 | 37.867 | 14.883 |
| ARPE-19 | 0.615 | 2.855 | 4.299 |
| SKOV-3 | 0.887 | 2.327 | 2.624 |
| Hep G2 | 1.802 | 3.31 | 1.836 |
| HT-29 | 1.031 | 1.777 | 2.27 |
| A549 | 0.665 | 1.534 | 2.304 |
| JKT-1 | 1.9 | 3.08 | 1.613 |
| T-47D | 6.078 | 13.811 | 2.272 |
| MCF-7 | 2.633 | 7.56 | 2.87 |
| MDA-MB-231 | 1.063 | 1.338 | 1.257 |
| HaCat | 2.639 | 4.205 | 1.593 |
| A431 | — | 4.042 | — |
| WM793 | 2.05 | 5.275 | 2.572 |
| T24 | 0.455 | 0.849 | 1.866 |
| 5637 | 1.817 | 5.477 | 3.013 |

**Table S3.**
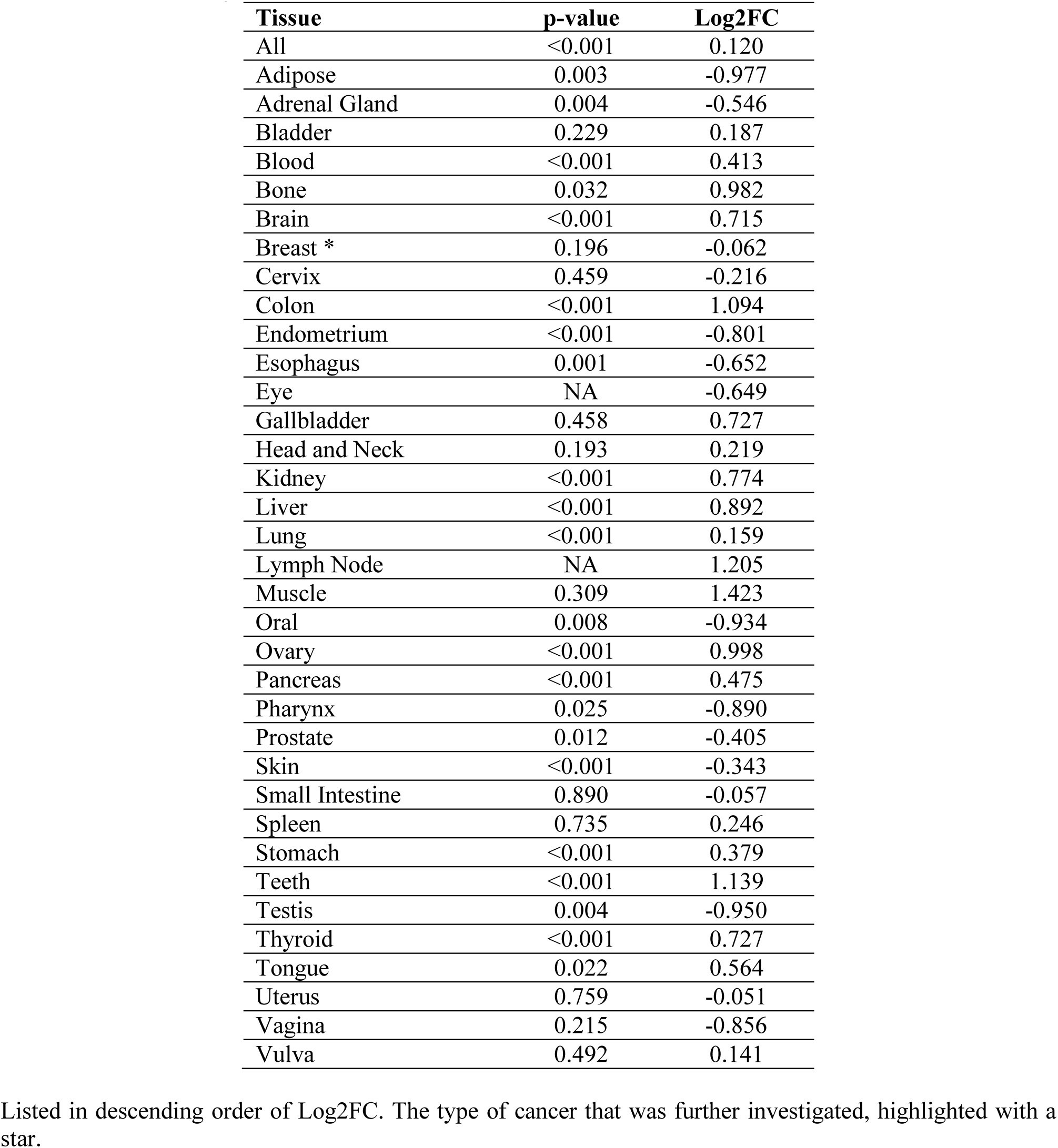
Statistical analysis of *TKS4* gene expression in different tumor types based on the GENT2 database, related to Figures 3A, S1.

**Table S4.**
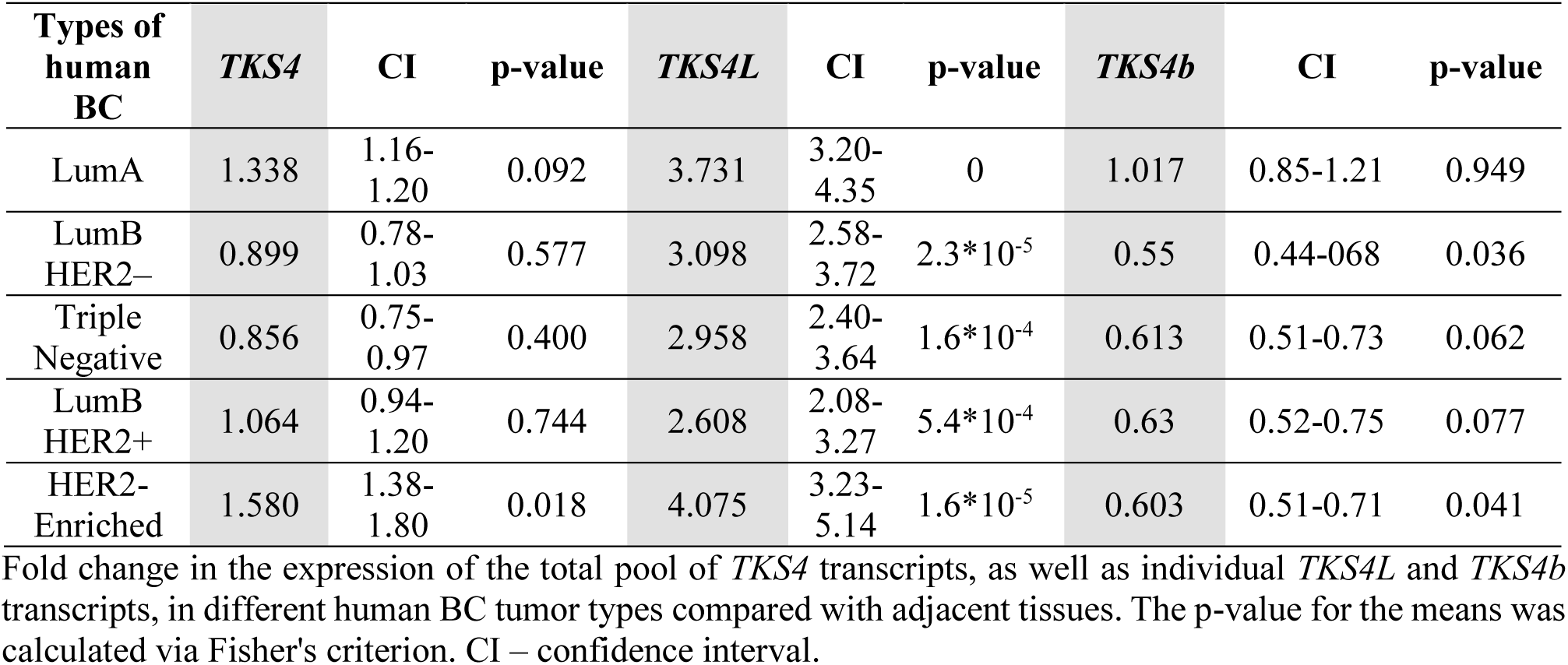
Expression of the *TKS4* gene in BC tumors, related to Figures 3D.

| Types of human BC | *TKS4* | CI | p-value | *TKS4L* | CI | p-value | *TKS4b* | CI | p-value |
| --- | --- | --- | --- | --- | --- | --- | --- | --- | --- |
| LumA | 1.338 | 1.16-1.20 | 0.092 | 3.731 | 3.20-4.35 | 0 | 1.017 | 0.85-1.21 | 0.949 |
| LumB HER2– | 0.899 | 0.78-1.03 | 0.577 | 3.098 | 2.58-3.72 | $2.3 \times 10^{-5}$ | 0.55 | 0.44-0.68 | 0.036 |
| Triple Negative | 0.856 | 0.75-0.97 | 0.400 | 2.958 | 2.40-3.64 | $1.6 \times 10^{-4}$ | 0.613 | 0.51-0.73 | 0.062 |
| LumB HER2+ | 1.064 | 0.94-1.20 | 0.744 | 2.608 | 2.08-3.27 | $5.4 \times 10^{-4}$ | 0.63 | 0.52-0.75 | 0.077 |
| HER2-Enriched | 1.580 | 1.38-1.80 | 0.018 | 4.075 | 3.23-5.14 | $1.6 \times 10^{-5}$ | 0.603 | 0.51-0.71 | 0.041 |

Fold change in the expression of the total pool of *TKS4* transcripts, as well as individual *TKS4L* and *TKS4b* transcripts, in different human BC tumor types compared with adjacent tissues. The p-value for the means was calculated via Fisher's criterion. CI – confidence interval.

**Table S5.** Ratios of *TKS4* transcripts expression in human BC tumor types and adjacent tissues, related to Figures 3E.

| Types of human BC | *TKS4L/TKS4b* | *TKS4b/TKS4L* |
| --- | --- | --- |
| Adjacent | 0.21 | 4.76 |
| LumA | 0.67 | 1.48 |
| LumB HER2– | 1.2 | 0.84 |
| Triple Negative | 0.89 | 1.13 |
| LumB HER2+ | 0.78 | 1.27 |
| HER2-Enriched | 1.28 | 0.78 |

**Table S6.** Genes expression in TKS4-cell lines, related to Figures 6C-D.

| mRNA name | MCF-7ΔTKS4 | MCF-7+TKS4L | MCF-7+TKS4b |
| --- | --- | --- | --- |
| *TKS4L* | 0.29*** | 137.937*** | 0.578*** |
| *TKS4b* | 0.249*** | 0.3*** | 8693.201*** |
| *TKS4d* | 0.219*** | 1.03 | 0.722 |
| *TKS5* | 0.000018*** | 0.649** | 0.603** |
| *TKS5L* | 0.048*** | 0.257*** | 0.407*** |
| *WIP* | 0.739** | 0.241*** | 0.473*** |
| *WASL* | 0.197*** | 0.033*** | 0.163*** |
| *CR16* | 0.075*** | 0.272*** | 0.678** |
| *TTP* | 2.711** | 0.309*** | 0.4** |

Fold change in the expression of *TKS4L*, *TKS4b*, *TKS4d*, *TKS5*, *TKS5L*, *WIP*, *WASL*, *CR16* and *TTP* transcripts in MCF-7ΔTKS4, MCF-7+TKS4L, and MCF-7+TKS4b cells compared with that in MCF-7wt cells. The p-value for the means was calculated via Fisher's criterion. \* – $p < 0.05$ , \*\* – $p < 0.01$ , \*\*\* – $p < 0.001$

**Table S7.**
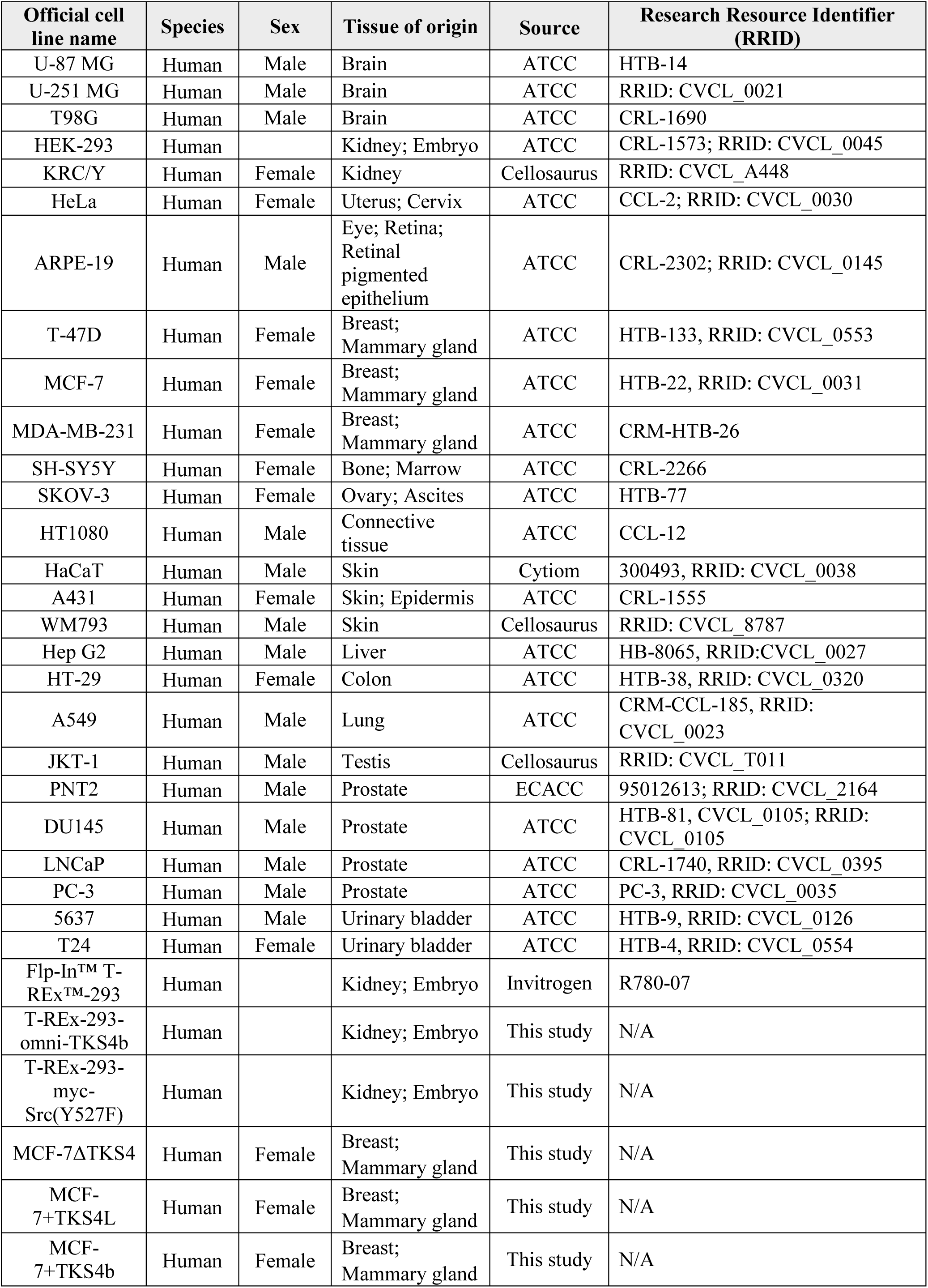
Cell lines. Related to METHODS «Cell culture and transfection».

**Table S8.** Antibodies. Related to METHODS «Antibodies and immunofluorescence reagents».

| **Antibodies** | **Source** | **Research Resource Identifier (RRID)** |
| --- | --- | --- |
| Anti-omni (D-8) mouse | Santa Cruz | Cat# sc-7270; RRID: AB_675763 |
| Anti-omni (M-21) rabbit | Santa Cruz | Cat# sc-499; RRID: AB_675765 |
| Anti-Vimentin (V9) mouse | Santa Cruz | Cat# sc-6260; RRID: AB_628437 |
| Anti-SMA (Smooth Muscle Actin) (CGA7) mouse | Santa Cruz | Cat# sc-53015; RRID: AB_628683 |
| Anti-myc (9E10) mouse | Santa Cruz | Cat# sc-40; RRID: AB_627268 |
| Anti-phosphotyrosine (PY99) mouse | Santa Cruz | Cat# sc-7020 |
| Anti-flag M2 mouse | Sigma Aldrich | Cat# F1804; RRID: AB_262044 |
| Anti-β-actin mouse | Sigma Aldrich | Cat# A1978; RRID: AB_476692 |
| Anti-α-tubulin mouse | Sigma Aldrich | Clone B-5-1-2; Cat# T6074 |
| Anti-SH3PXD2B rabbit | Sigma Aldrich | Cat# HPA058141 |
| Anti-GST goat | Sigma Aldrich | Cat# GE27-4577-01 |
| Anti-N-Cadherin rabbit | Abcam | Cat# ab18203; RRID: AB_444317 |
| Anti-E-Cadherin rabbit | Abcam | Cat# ab40772; RRID: AB_731493 |
| Anti-CTTN (p80/85) mouse | Sigma Aldrich | Clone 4F11; Cat# 05-180-I |
| Goat anti-rabbit IgG, HRP Conjugate | Promega | Cat# W4011; RRID: AB_430833 |
| Goat anti-mouse IgG, HRP Conjugate | Promega | Cat# W4021; RRID: AB_430834 |
| Donkey anti-goat IgG, HRP Conjugate | Promega | Cat# V8051; RRID: AB_430838 |
| Goat anti-mouse Alexa Fluor 633 | Thermo Fisher | Cat# A-21050; RRID: AB_2535718 |
| Donkey anti-rabbit Alexa Fluor 555 | Abcam | Cat# ab150074; RRID: AB_2636997 |
| Goat anti-mouse Alexa Fluor 488 | Invitrogen | Cat# A-11001; RRID: AB_2534069 |
| Alexa Fluor 555-phalloidin | Invitrogen | Cat# A34055 |

**Table S9.**
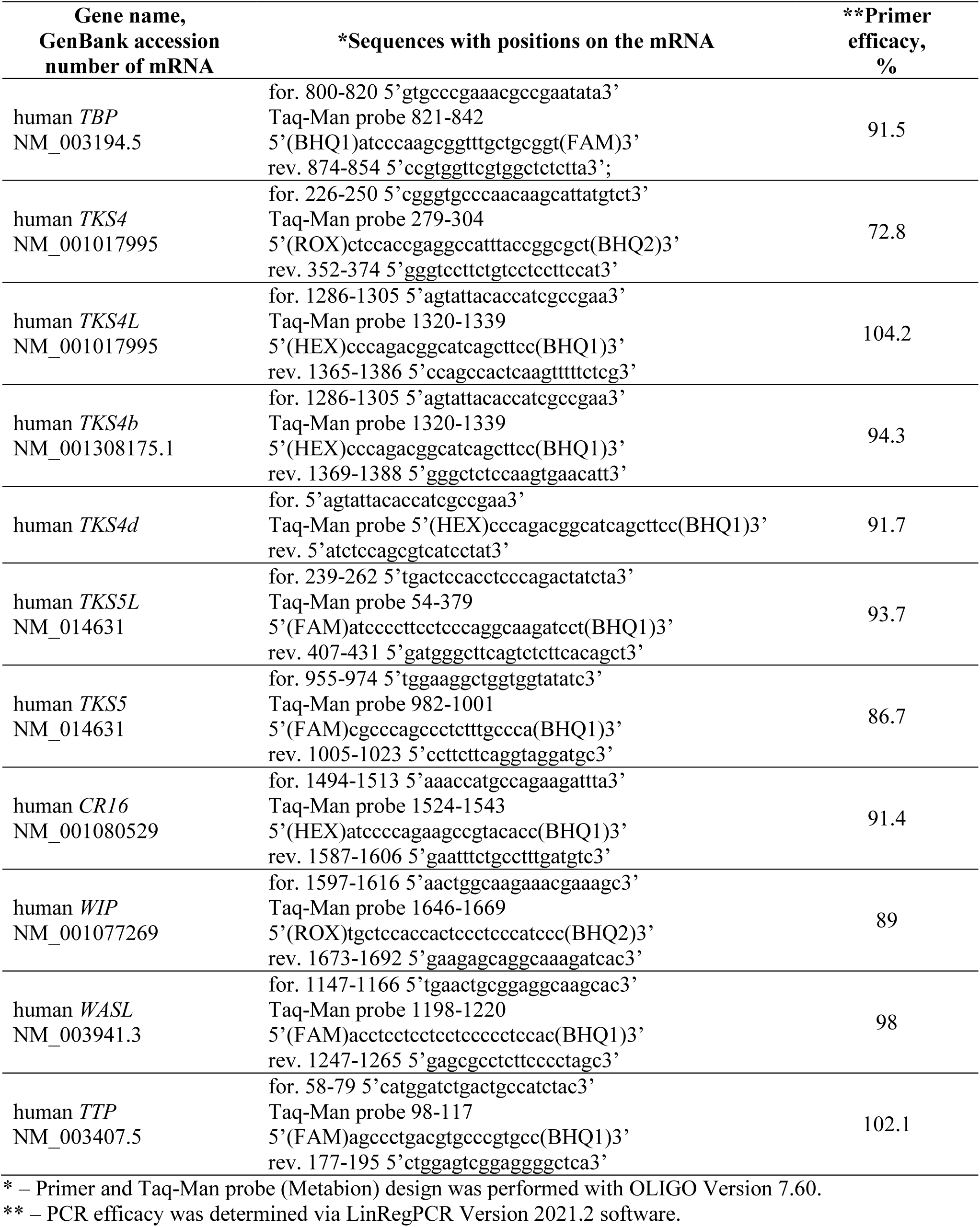
Primers and Taq-Man probes used for qPCR. Related to METHODS «cDNA synthesis and qPCR with fluorescence labeled probes».

---

**Figure S1.**
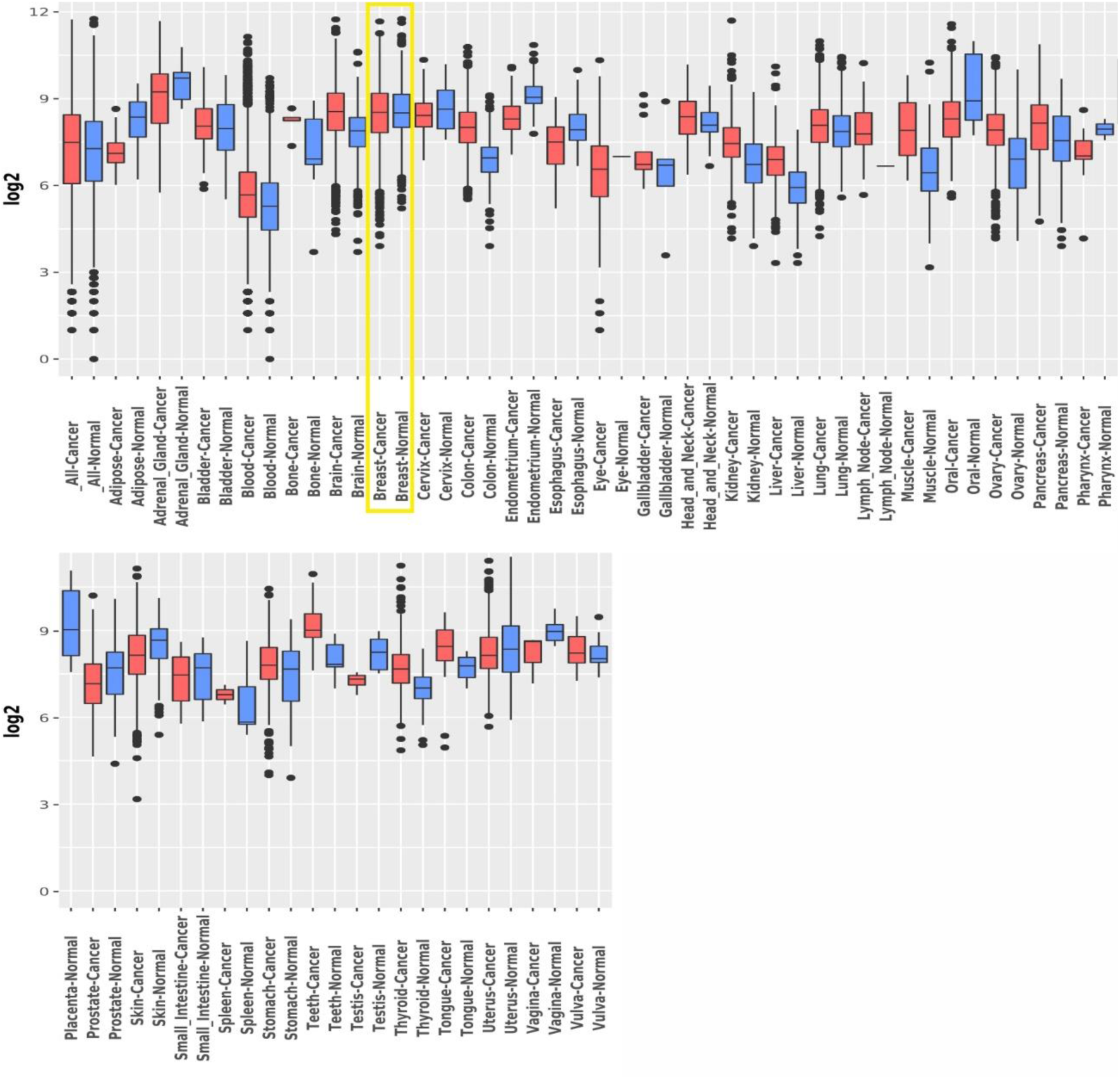
*TKS4* expression in normal and tumor samples in various cancer types based on the GENT2 database – full version, related to Figures 3A. The BC is highlighted with a box.

**Figure S2.**
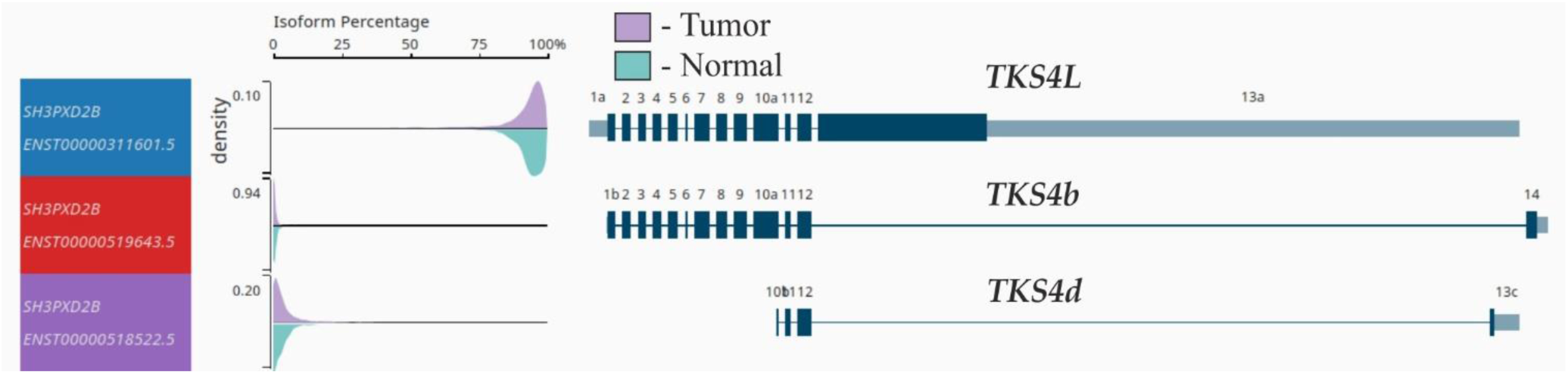
Expression of the *TKS4L*, *TKS4b*, and *TKS4d* transcripts in normal and breast tumor tissues based on analysis of the TCGA dataset using the UCSC Xena browser, related to RESULTS «Expression profile of *TKS4* transcripts in breast tumors».

**Figure S3.**
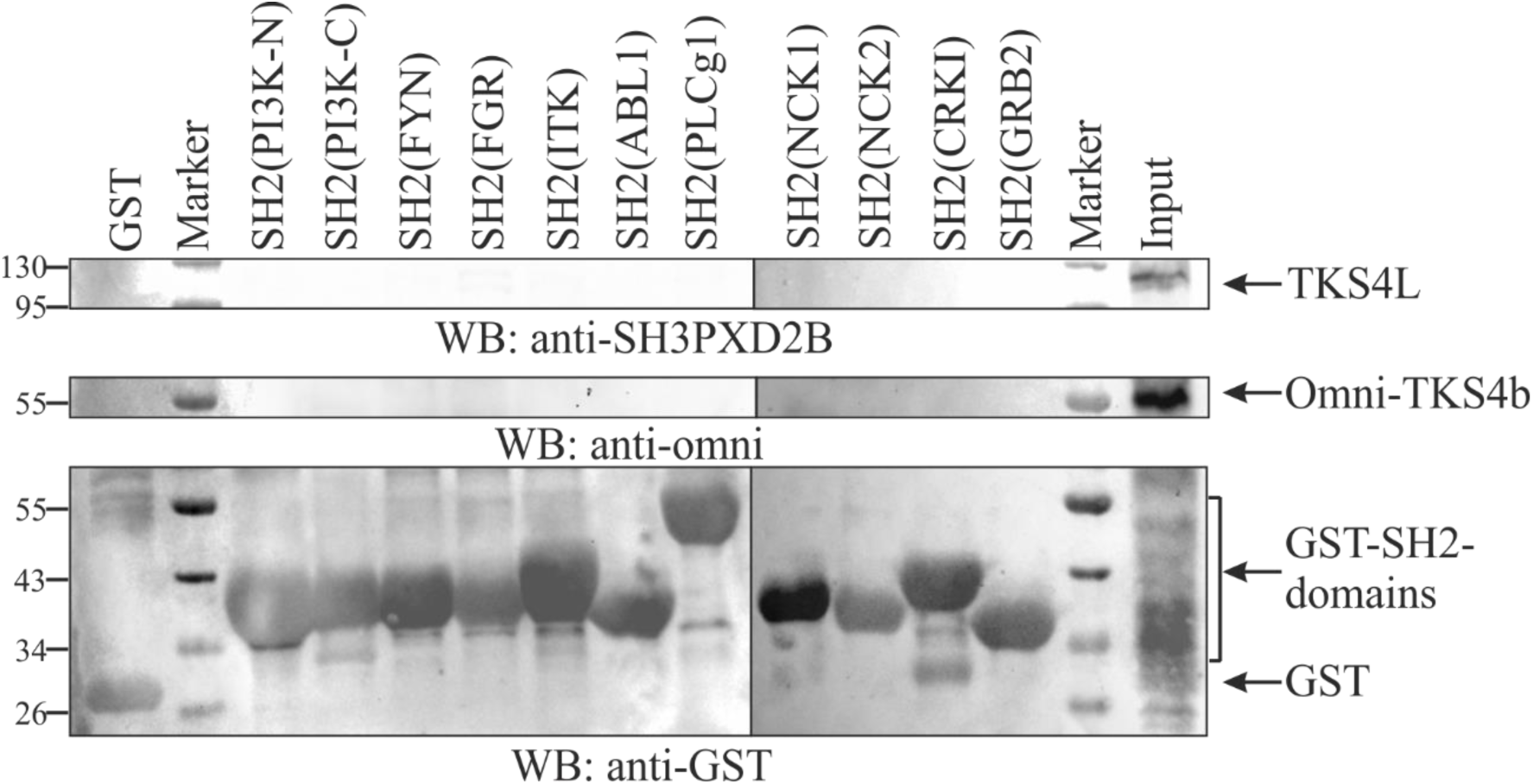
**The TKS4 isoforms are not recognized by the SH2 domains without overexpression Src(Y527F)**, related to Figures 5B. Analysis of TKS4L and TKS4b isoform interactions with the GST-SН2 domains of PI3K, FYN, FGR, ITK, ABL, PLCg1, NCK1, NCK2, CRKI, GRB2 proteins and GST alone (control), which were immobilized on glutathione beads and incubated with lysates from growing T-REx-293 cells with transiently expressed omni-TKS4b. Bound proteins were analyzed by Western blotting using antibodies against TKS4L, anti-omni and anti-GST antibodies. WB, Western blot.

**Figure S4.**
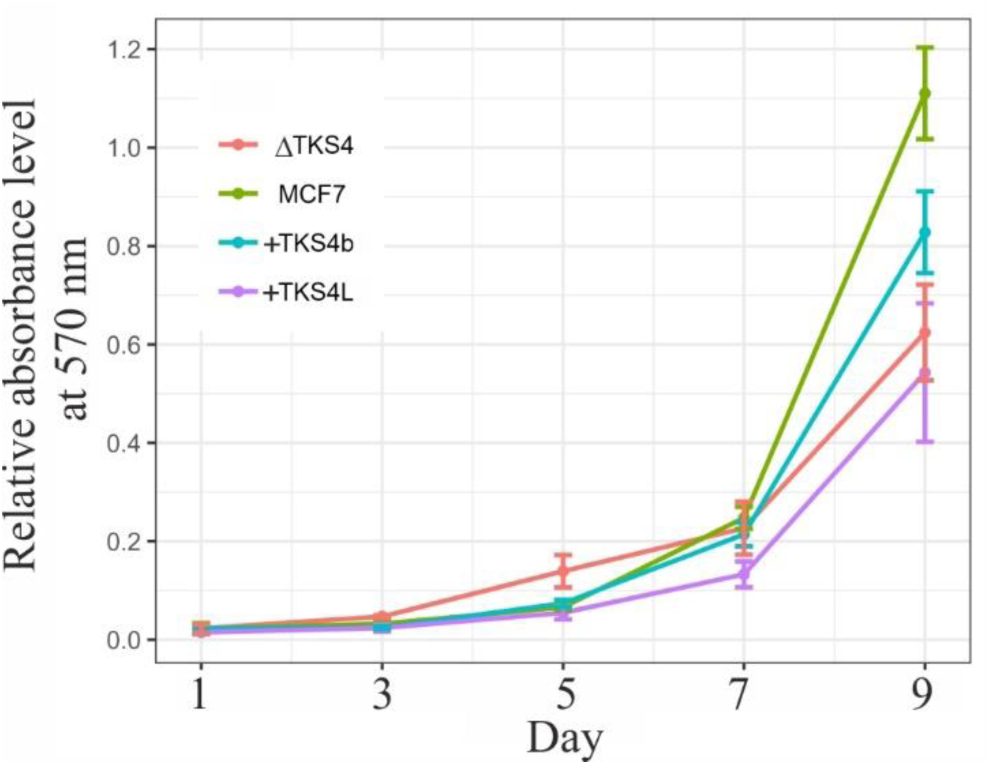
**Proliferation rates of the cell lines MCF-7wt, MCF-7ΔTKS4, MCF-7+TKS4L, and MCF-7+TKS4b (M±σ, n=6)**, related to Table 1.

**Figure S5.**
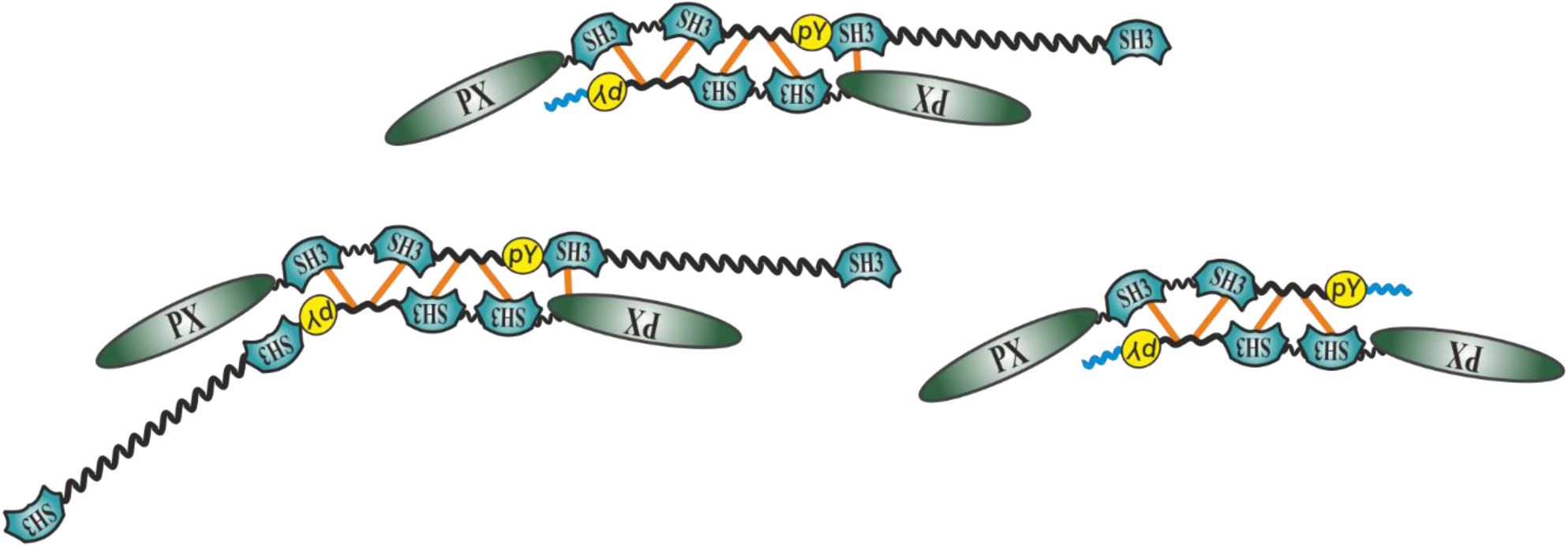
**Hypothetical interaction of the TKS4L and TKS4b isoforms after phosphorylation**, related to Discussion. Hypothetical schemes for the formation of homo-and heterodimers of the TKS4L and TKS4b isoforms after phosphorylation, assuming interaction between the SH3_1 and SH3_2 domains and the TAM motif of another molecule.

**Figure S6.**
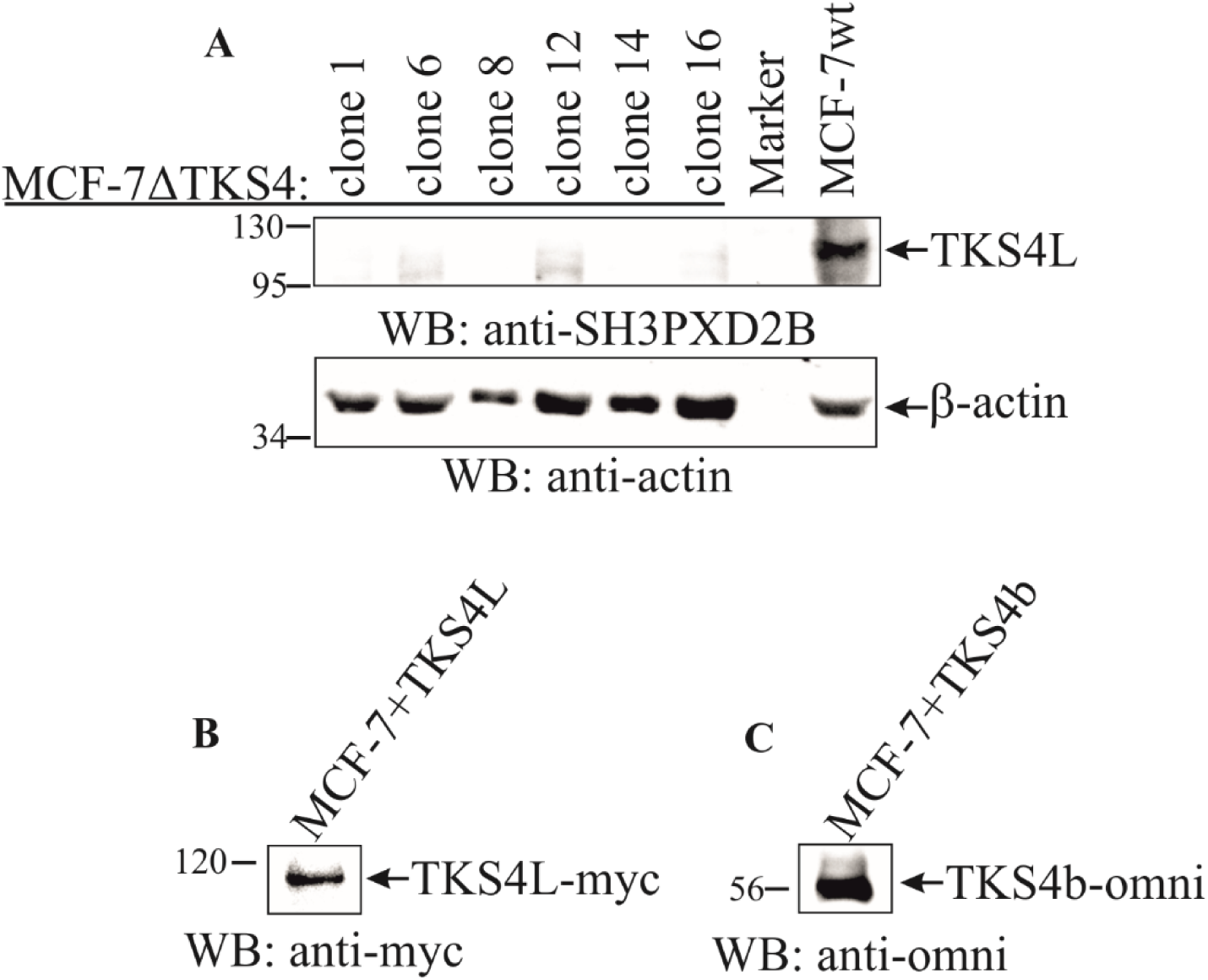
Western blot analysis of the MCF-7 cell lines with knockout and overexpression of TKS4. Related to METHODS «Generation of MCF-7 stable cell lines with a knockout and overexpression of *TKS4* gene». **(A)** WB analysis of *TKS4* gene knockout in MCF-7ΔTKS4 cell clones. *TKS4* gene was edited with CRISPR/Cas9 technology. SH3PXD2B antibodies against the C-terminal region of TKS4 were used for the analysis. β-Actin was used as a loading control; **(B)** WB analysis of TKS4L-myc overexpression in MCF-7+TKS4L cell line; **(C)** WB analysis of TKS4b-omni overexpression in MCF-7+TKS4b cell line. TKS4L-myc and TKS4b-omni overexpression was performed with pcDNA4/TO methodology.

